# Regulation of Harvester Ant Foraging as a Closed-Loop Excitable System

**DOI:** 10.1101/322974

**Authors:** Renato Pagliara, Deborah M. Gordon, Naomi Ehrich Leonard

**Affiliations:** Department of Mechanical and Aerospace Engineering, Princeton University, Princeton, New Jersey, United States of America; Department of Biology, Stanford University, Stanford, California, United States of America

## Abstract

Ant colonies regulate activity in response to changing conditions without using centralized control. Harvester ant colonies forage in the desert for seeds, and their regulation of foraging manages a tradeoff between spending and obtaining water. Foragers lose water while outside in the dry air, but the colony obtains water by metabolizing the fats in the seeds they eat. Previous work shows that the rate at which an outgoing forager leaves the nest depends on its recent experience of brief antennal contact with returning foragers that carry a seed. We examine how this process can yield foraging rates that are robust to uncertainty and responsive to temperature and humidity across minutes to hour-long timescales. To explore possible mechanisms, we develop a low-dimensional analytical model with a small number of parameters that captures observed foraging behavior. The model uses excitability dynamics to represent response to interactions inside the nest and a random delay distribution to represent foraging time outside the nest. We show how feedback of outgoing foragers returning to the nest stabilizes the incoming and outgoing foraging rates to a common value determined by the “volatility” of available foragers. The model exhibits a critical volatility above which there is sustained foraging at a constant rate and below which there is cessation of foraging. To explain how the foraging rates of colonies adjust to temperature and humidity, we propose a mechanism that relies on foragers modifying their volatility after they leave the nest and get exposed to the environment. Our study highlights the importance of feedback in the regulation of foraging activity and points to modulation of volatility as a key to explaining differences in foraging activity in response to conditions and across colonies. Our results present opportunities for generalization to other contexts and systems with excitability and feedback across multiple timescales.

**Author Summary:** We investigate the collective behavior that allows colonies of desert harvester ants to regulate foraging activity in response to environmental conditions. We develop an analytical model connecting three processes: 1) the interactions between foragers returning to the nest and available foragers waiting inside the nest, 2) the effect of these interactions on the likelihood of available foragers to leave the nest to forage, and 3) the return of foragers to the nest after finding seeds. We propose a mechanism in which available foragers modify their response to interactions after their first exposure to the environment. We show how this leads to colony foraging rates that adjust to environmental conditions over time scales from minutes to hours. Our model may prove useful for studying resilience in other classes of systems with excitatory dynamics.

## Introduction

Social insect colonies operate without central control. Colonies maintain coherence and plasticity in the face of perturbation and change, even though individuals have limited and uncertain information on the state of the group and the state of the environment. Collective behaviors are understood to emerge from response of individuals to social interactions and measurements of the local environment [1–4]. The study of social insects provides opportunities to investigate open, fundamental questions on how collective behavior adjusts to different conditions and how small differences in these adjustments can lead to large differences in behavior across groups.

The regulation of foraging activity in colonies of the harvester ant (*Pogonomyrmex Barbatus*) is a well-studied example of collective behavior [5]. Harvester ants live in the hot and dry Southwestern US desert where they forage for seeds scattered by wind and flooding. Foragers do not use pheromone trails; instead, they spread out across selected foraging areas in search of seeds [6]. The regulation of foraging activity manages a tradeoff between spending and obtaining water: foragers lose water while outside in the dry air, but colonies obtain water by metabolizing the fats in the seeds that they eat [7,8].

Harvester ant colonies regulate the rate at which foragers leave the nest using the incoming rate of successful foragers returning with food [9–13]. When an ant contacts another ant with its antennae, it perceives the other ant’s cuticular hydrocarbon (CHC) profile [9]. Because conditions outside the nest change the chemistry of the cuticular hydrocarbons, CHC profiles are task-specific [14]. In the course of antennal contact, one ant can detect whether another is a forager. An available forager, waiting in the entrance chamber inside the nest, is stimulated to leave the nest by antennal contact with foragers carrying food [11–13]. Because each forager searches until it finds a seed, the rate of interaction serves as a noisy measurement of the current foraging conditions [6,15]. A higher rate of forager return, which reflects a greater food supply, increases the likelihood that available foragers will leave the nest to forage [12,13,16].

In the integrator model of [13], each available forager inside the nest collects evidence from incoming foragers by integrating its recent experience of antennal contacts. When the integrated stimulus passes a threshold, the available forager is likely to leave the nest; in the absence of interactions the forager is likely to descend from the entrance chamber to the deeper nest [12,16], protecting the colony from the inherently noisy signal that results from limited and uncertain interactions [17]. The integrator model has been used to study regulation of the outgoing foraging rate on short timescales [15].

Colonies regulate their foraging activity on longer timescales, such as from hour to hour, from day to day [18,19], and across years [5,18,20,21] as colonies grow older and larger. Over timescales from tens of minutes to hours, ants that start as available foragers inside the nest leave the nest to forage, find seeds, return to the nest, and become available foragers again. Thus, the activation of available foragers inside the nest through interactions with incoming foragers is connected in a “closed loop” to the foraging activity outside the nest through feedback of the ants themselves: the stream of foraging ants out of the nest is the input to the foraging activity, and the output of the foraging activity is the stream of foraging ants into the nest (see Fig. 1). However, little is known about the role of feedback in the regulation of foraging activity at the timescale of hours and as foraging activity is adjusted to changing environmental conditions. By mid-day in the summer, temperature is high and humidity is low (Fig.S1). Foraging activity increases from its start in early morning, often levels off and declines to no activity during the heat of the afternoon.

**Fig. 1.**
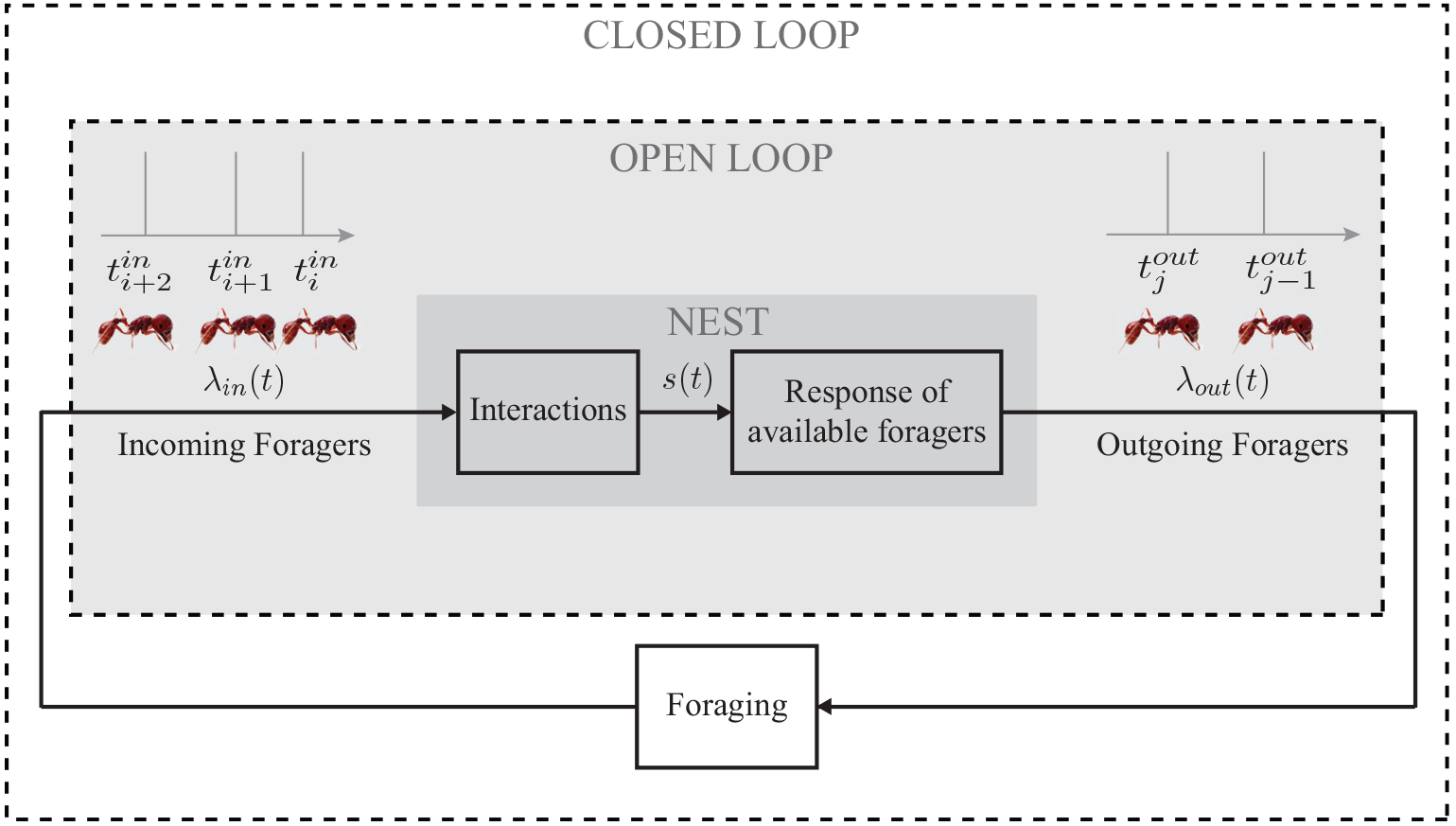
Diagram of the closed-loop model with two components inside the nest and one component outside the nest. The “Interactions” component maps the sequence of incoming foragers λ_*in*_ to a stimulus *s* to represent the result of interactions of available foragers inside the nest entrance chamber with returning food-bearing foragers. The mapping uses a leaky integrator that increases by a fixed magnitude with every incoming forager and has a natural decay rate. The “Response of available foragers” component maps *s* to the sequence of outgoing foragers λ_*out*_ using the nonlinear FN oscillator dynamics. Each oscillation represents an ant leaving the nest to forage. The “Foraging” component maps λ_*out*_ to λ_*in*_ using a random time delay with an associated probability distribution to represent the time an ant spends outside the nest foraging.

How a colony regulates foraging in response to environmental conditions, especially temperature and humidity, is ecologically important. Colonies live for 20-30 years. At about five years of age a colony begins to produce reproductives that mate and found offspring colonies [22]. Colonies differ in the regulation of foraging and these differences persist from year to year, including variation in how often colonies are active [18] and in how they respond to changing temperature and humidity conditions [5,19,21]. How a colony adjusts foraging activity to low humidity and high temperature is crucial for reproductive success: colonies that conserve water are more likely to have offspring colonies [5]. Here we model how colonies adjust to environmental conditions, to provide insight into the ecologically important variation among colonies that underlies the evolution of collective behavior.

Previous modeling work has elucidated how the outgoing foraging rate depends on the incoming foraging rate [15], and how individuals assess interaction rate [13]. But we do not know how these are combined to adjust foraging activity across minutes to hours-long timescales, how the adjustments may depend on environmental conditions, or how they may differ from colony to colony.

Here we propose a closed-loop model (Fig. 1) to address these questions by examining how a returning forager’s assessment of external conditions provides additional feedback to the colony and in turn adjusts the colony foraging rate. Our model is motivated in part by the frequent use of excitability dynamics to model neurons, and the parallels between ant-to-ant interactions that drive foraging and neuron-to-neuron interactions that underlie the cognitive abilities of organisms [13,23–24,25 26]. Using well-studied excitability dynamics of a weakly interacting collective, we introduce feedback at multiple time-scales and explore general questions concerning stability and responsiveness to a changing environment.

Drawing on theory and tools from dynamical and control systems, we study how the incoming and outgoing foraging rates adjust with time at timescales much longer than previously considered [13,15]. We ask how the rates can stably approach a common value even in the presence of disturbance, how adjustments and stability of rates differ under various foraging conditions, and how sensitivities to parameters may explain differences in foraging behavior across colonies.

## Methods

### Field Observations of Foraging Activity

We performed field observations of red harvester ant colonies at the site of a long term study near Rodeo, New Mexico, USA. Observations were made in August and September of 2015, 2016, and 2017. Foragers leave the nest in streams or trails that can extend up to 20 m from the nest [27]. Each forager leaves the trail to search for seeds, and once it finds food, it returns to the nest [6,27]. Data on foraging rates were recorded from the beginning of the foraging period in early morning until around noon. We recorded the times at which foragers crossed a line perpendicular to the trail at a distance of about 1 m from the nest entrance, as in previous work (e.g. [15,21,28]). The timestamps for each forager crossing the line were recorded either manually in real-time with the assistance of an electronic tablet and custom software, or from video recordings, processed with computer vision software (AnTracks Computer Vision Systems, Mountain View, CA). In some cases we used both tablet and video to ensure that both data collection methods provided similar results.

We denote by 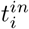, *i* ∈ ℕ, the sequence of times incoming foragers cross the line and by 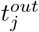, *j* ∈ ℕ, the sequence of times outgoing foragers cross the line. Sequences of incoming and outgoing foragers are represented as sums of infinitesimally narrow, idealized spikes in the form of Dirac-delta functions:

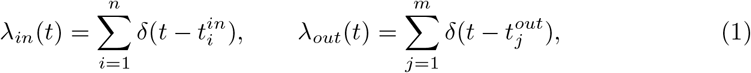

where *n* and *m* are the indices of the last incoming and outgoing forager, respectively, before time *t*. We estimated the instantaneous incoming and outgoing foraging rates, in units of ants/sec, using a sliding window filter with window Δ*t* = 300 sec:

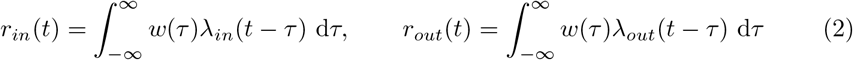

where

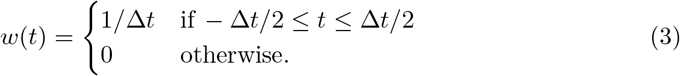

### Model

We propose a low-dimensional model with a small number of parameters that has sufficiently rich dynamics to capture the range of observed foraging behavior across minute to hour-long timescales and yet retains tractability for analysis. We use the model to systematically investigate the effects of model parameters and environmental conditions, notably temperature and humidity, on foraging rates.

Our model has three components as shown in Fig. 1: 1) the *Interactions* component models the accumulation of evidence by available foragers inside the nest entrance chamber from their interactions with incoming foragers carrying food, 2) the *Response of available foragers* component models the activation of available foragers to leave the nest to forage in response to accumulated evidence, and 3) the *Foraging* component models the collecting of seeds outside the nest by active foragers. We assume the total number of foragers *N* (active foragers outside the nest plus available foragers inside the nest) remains constant throughout the foraging day, although this assumption can be relaxed in a generalization of the model.

#### Interactions

We use leaky-integrator dynamics to model the stimuli *s* that the group of available foragers inside the nest entrance chamber experience from their interactions with returning food-bearing foragers:

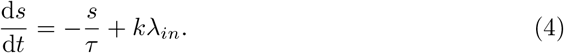

The continuous-time signal *s* increases by a fixed amount *k* with every incoming forager in λ_*in*_ and decays exponentially back to zero with a time constant of τ.

The leaky-integrator dynamics work as an evidence accumulator that gradually forgets past evidence. These dynamics have been used to model chemical synapses [29] and have been used as the integrate-and-fire neuronal model when there is no reset boundary [30–32].

#### Response of Available Foragers

We first consider a homogeneous colony and model the scalar activation state *υ* of available foragers in the nest entrance chamber as the fast timescale variable in the FitzHugh-Nagumo (FN) equations [33,34] often used to model neuronal excitability:

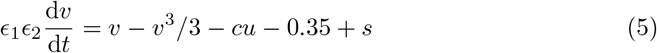

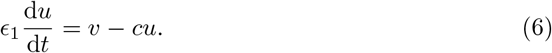

The FN equations describe nonlinear oscillator dynamics with the stimulus *s* of Eq. (4) as the input. Oscillations result from a balance between positive feedback in *υ* (first term on the right of Eq. (5)) and negative feedback in the slow timescale variable *u*. We refer to the parameter *c*, which scales the negative feedback and modulates the frequency of oscillations, as the *volatility* of the available foragers. The parameter *ϵ*_2_ defines the time separation between the dynamics of the fast and slow states, and the parameter *ϵ*_1_ defines the time separation between the FN dynamics and the stimulus dynamics (4). The constant 0.35 is chosen so that the threshold input required to elicit oscillations is lower than *k*, the magnitude by which *s* increases with every incoming forager.

The activation dynamics (Eqs. (5) and (6)) of the available foragers yield three qualitatively distinct dynamical regimes, determined by the magnitude of input *s*. In the first regime, the system remains in a *resting* state for low values of *s*. This reflects the situation in which there are In the second regime, which takes place when *s* ≥ *b*_1_ > 0, the system is in an excited state with relaxation oscillations in *υ*. This reflects the situation in which incoming foragers are sufficiently frequent to stimulate the available foragers. The transition from resting to oscillatory behavior as *s* increases corresponds to a Hopf bifurcation and *b*_1_ is the corresponding bifurcation point. The oscillations appear as short-lived spikes, and we define each spike for which *υ* increases above 0.75 as a forager leaving the nest. The shortest possible time between foragers leaving the nest is determined by the volatility *c* (see SI Appendix 1).

In the third regime, corresponding to very large values of *s* ≥ *b*_2_ > *b*_1_, there are no oscillations and the system is fixed in a *saturation* state. The transition from oscillatory to saturated regime is a second Hopf bifurcation with bifurcation point *b*_2_. We associate the saturated state of the FN model with conditions under which high instantaneous incoming rates lead to a decrease in the instantaneous outgoing rate. These conditions include 1) overcrowding effects, which reduce the percentage of interactions experienced by each available forager relative to the incoming foraging rate, 2) the limited size of the nest entrance tunnel, which constrains how many foragers can enter and leave the nest in a short amount of time, and 3) the difference in timescales between the high outgoing rates, in seconds, and the time required, in minutes, for foragers to move from the deeper chambers of the nest up to the entrance chamber [12,16].

#### Foraging

We treat the process of foraging for seeds outside the nest as a random time delay. We model the interval between the time that a forager leaves the nest and the time when it returns with food as a chi-square random variable *X*, with parameter *D* representing the mean foraging time in minutes. The distribution of foraging times *F*(*X*, *D*) is

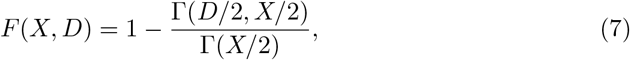

where Γ(*X*) and Γ(*a*, *X*) are the Gamma function and the upper incomplete Gamma function, respectively. This right-skewed distribution is based on field observations of the duration of foraging trips, measured as the total time elapsed from when a forager leaves the nest to when it returns with food [6]. For *D* = 2, *F*(*X*, 2) = 1 – *e*^−*X*/2^.

Our model for the foraging process is equivalent to a queueing system [35] in which arriving customers, represented by outgoing foragers λ_*out*_, that find a seed after a given random service time. The number of servers in the foraging process queue is assumed to be infinite because foragers do not need to wait before they start looking for a seed (i.e., before receiving the service).

#### Proposed Mechanism for Response to Environmental Conditions

We propose a mechanism for colony response to environmental conditions, illustrated in Fig. 2, in which the volatility of a forager changes after it has been on a foraging trip and exposed to the conditions outside the nest. The proposed mechanism is based on measurements showing that the temperature and humidity inside the nest remain constant throughout the foraging activity period (see Fig. S1). This means that foragers have no information about conditions outside until they leave the nest.

As a first approximation, the model changes the volatility of each forager after it leaves the nest to forage for the first time. Available foragers that have not yet been outside the nest, and are therefore uninformed about the current temperature and humidity outside the nest, have volatility *c*_*u*_. Available foragers that have been outside at least once to forage, and are therefore informed about the current temperature and humidity, have volatility *c*_*i*_. The values of *c*_*u*_ and *c*_*i*_ can be any positive real numbers, and these values, which represent an average uninformed and an average informed available forager in the colony, respectively, can vary across colonies and across days. Further, *c*_*i*_ is introduced explicitly to depend on conditions such as humidity and temperature outside the nest. For example, the hotter and drier it is outside, the smaller the *c*_*i*_, and the foragers become less volatile and thus less likely to make subsequent foraging trips. The cooler and more humid it is outside, the larger the *c*_*i*_, and the foragers become more volatile and thus more likely to make subsequent foraging trips.

**Fig. 2.**
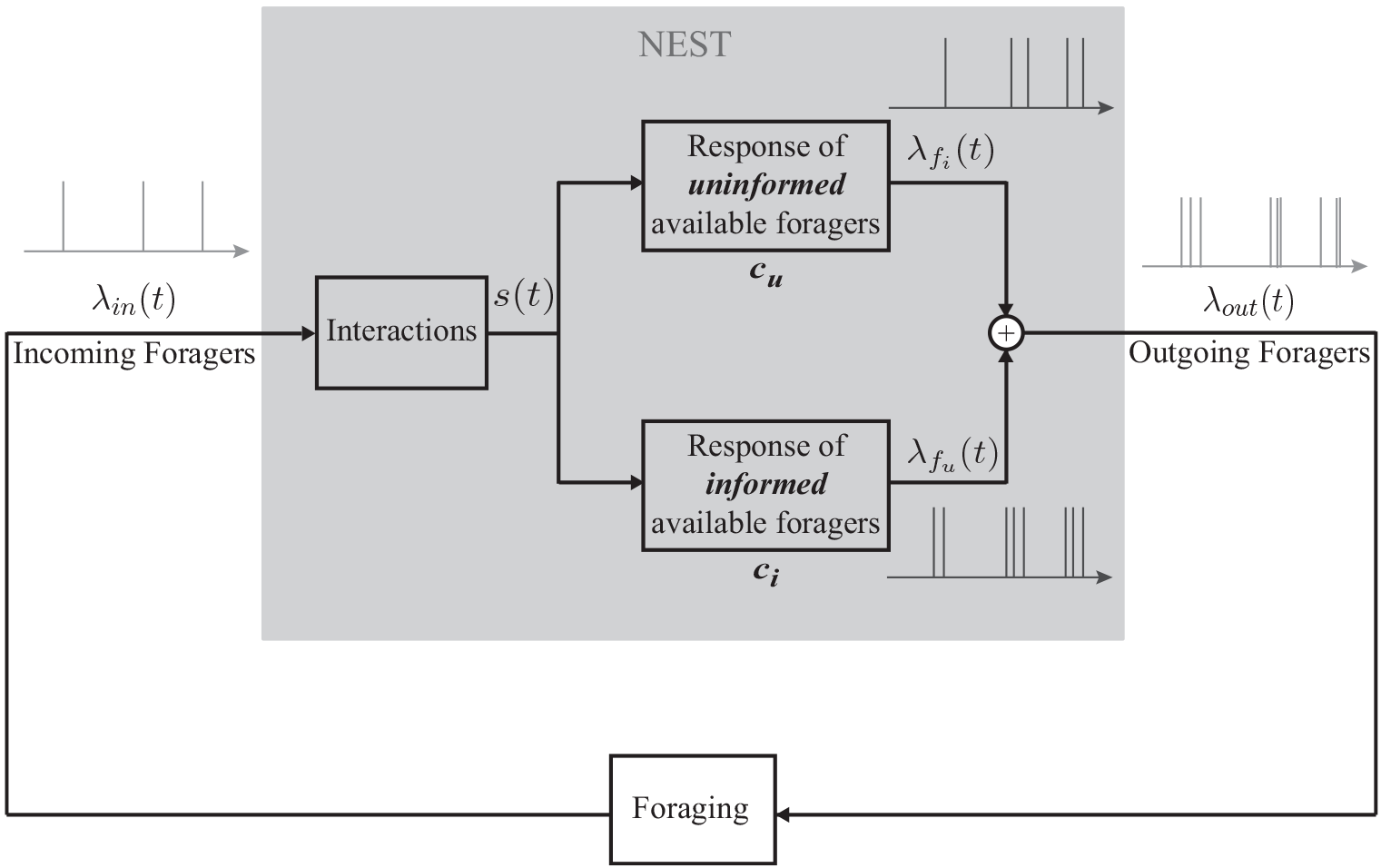
Block diagram of proposed mechanism for response of colony to environmental conditions. The available foragers inside the nest comprise two sets: *f*_*u*_ corresponds to those that have not yet left the nest and so are uninformed about the conditions outside the nest, and *f*_*i*_ corresponds to those informed during a previous foraging trip. The response of each set to *s* is represented by a different FN model, distinguished by the volatility parameter *c*_*u*_ for the uninformed and *c*_*i*_ for the informed. The outputs of these two oscillator dynamics are weighted probabilistically using thinning to get an outgoing stream of foragers λ_*out*_(*t*).

Let *f*_*u*_ be the set of *n*_*u*_ *uninformed* available foragers that have not yet left the nest during the day and thus have no information about the environmental conditions and *f*_*i*_ the set of *n*_*i*_ *informed* available foragers that have been exposed to the environmental conditions during one or more previous foraging trips that day. We assume that once a forager becomes informed, it remains informed for the rest of the foraging day. The ants in *f*_*u*_ have volatility *c*_*u*_ and the ants in *f*_*i*_ have volatility *c*_*i*_. Let *x*_*u*_ = *n*_*u*_/(*n*_*u*_ + *n*_*i*_) and *x*_*i*_ = *n*_*i*_/(*n*_*u*_ + *n*_*i*_) be the fraction of available foragers that are uninformed and informed, respectively, where we assume that *n*_*u*_ + *n*_*i*_ > 0. Then *x*_*u*_ + *x*_*i*_ = 1.

Initially, *x*_*i*_ = 0 and the colony is completely uninformed (*x*_*u*_ = 1). As foragers return to the nest after their first trip, *x*_*i*_ begins to increase and can continue to increase until *x*_*i*_ = 1 (*x*_*u*_ = 0), when all *N* foragers have been outside the nest at least once. How many minutes (or hours) it takes for *x*_*i*_ to transition from 0 to 1 depends on *N*, *D*, and the changing foraging rates. To model the changing foraging rates, we use two sets of FN oscillator dynamics: one to represent the response to *s* of the uninformed ants in *f*_*u*_ with volatility *c*_*u*_ and a second to represent the response to *s* of informed ants in *f*_*i*_ with volatility *c*_*i*_. Let the corresponding sequences of output from the two oscillator dynamics be λ*f*_*i*_ and λ*f*_*u*_, respectively. We define the sequence of outgoing foragers λ_*out*_ as a probabilistic sum of λ_*i*_ and λ_*u*_, using a method called *thinning* [36]: Every event in λ_*i*_ is kept in λ_*out*_ with probability *x*_*i*_, and every event in λ_*u*_ is kept in λ_*out*_ with probability 1 – *x*_*i*_. When *x*_*i*_ = 0 the foraging rate is determined by *c*_*u*_, and when *x*_*i*_ = 1 the foraging rate is determined by *c*_*i*_. When 0 < *x*_*i*_ < 1, the effective *c* will be a nonlinear combination of *c*_*u*_ and *c*_*i*_. The higher the effective *c*, the higher the outgoing foraging rate.

Here foragers adjust their volatility only once after their first foraging trip outside. We find that even with this adjustment at first exposure, the model provides the range of foraging behavior observed. However, the model can be generalized and predictions refined by allowing for adjustments on subsequent foraging trips, and by allowing for other kinds of adjustments. For example, more than two sets of available foragers with different values of volatility can be used to model effects of repeated exposures to the environment, changing conditions on successive trips, or decay of information about the external environment over time. A decrease in *N* (total number of foragers outside and available inside the nest) can be used to model active foragers that return to the deeper nest after exposure to hot and dry outside conditions [12].

### Results

#### Observations of Regulation of Foraging in Red Harvester Ants

Observations of instantaneous foraging rates computed from the 2015, 2016, and 2017 data show that across colonies and days, the incoming and outgoing foraging rates *r*_*in*_(*t*) and *r*_*out*_(*t*), where *t* is time of day, undergo a transient early in the foraging period followed by an equilibration to a near-equal value, i.e., *r*_*in*_(*t*) ≈ *r*_*out*_(*t*), during the middle part of the foraging period.

The equilibration of the incoming and outgoing foraging rates to a near-equal value lasts for intervals from tens of minutes to several hours, and so we refer to it as a quasi steady-state (QSS). We show the data for two colonies in Fig. 3. We plot the incoming rate *r*_*in*_ (blue) and the outgoing rate *r*_*out*_ (red) computed from the data for Colony 1357 (Fig. 3A) and Colony 1317 (Fig. 3B) versus time of day on August 20, 2016. For Colony 1357, the rates equilibrate to a near-equal value early in the day, i.e., between 8:00 and 8:30 am. This is followed by a couple of dynamic adjustments, but then by 9:30 am until just before noon, when all the ants returned to the nest, the incoming and outgoing rates were very closely equilibrated at a QSS rate of around 0.25 ants/sec. Colony 1317 also was observed to reach a QSS. Its incoming and outgoing rates equilibrate to a near equal value shortly after 10:00 am, which lasts until just before noon, when all the ants return to the nest. Colonies vary greatly in foraging rate [21], and that was true of these two as well. For Colony 1317, the QSS rate is approximately 0. 65 ants/sec, more than twice the QSS rate for Colony 1357 on the same day.

We show data for two other colonies in Fig. 4. Fig. 4A and C show *r*_*in*_ (blue) and *r*_*out*_ (red) versus time of day for a single colony, Colony 664, on two different days: August 27, 2015 and August 31, 2015. In each plot, the rates can be seen to come to a near-equal value sometime after 10:30 am. We plot in green the difference between number of incoming and number of outgoing foragers versus the time of day. The rates are at a QSS when the green curve is approximately horizontal. These data show, as has been observed previously [37], that a given colony varies in foraging rate from day to day, demonstrating that foraging is regulated by processes other than the number of available foragers. From Fig. 4A and C it can be seen that Colony 664 reaches a QSS rate on August 27, 2015 that is more than twice the QSS rate it reaches on August 27, 2015. We note that August 27, 2015 was slightly cooler and more humid than August 31, 2015. On August 27 the average temperature and humidity were 25.3 C and 58% while on August 31 they were 26.8 C and 53%. Moreover, at 11am on August 27, they were 27.5 C and 52% while at 11am on August 31, they were 28.8 C and 45%. Fig. 4E shows the data for Colony 863 on September 1, 2015, which were recorded manually. No QSS is observed, i.e., the ants go out but then return to the nest by 11:00 am without maintaining a steady-state of foragers outside of the nest. Colony 863 did reach a QSS at a reasonably high foraging rate at 11:00 am on September 5, 2015 (see Fig. S2A). These observations are consistent with measurements showing that September 1, 2015 was much hotter and drier than September 5, 2015. On September 1 the average temperature and humidity were 25.2 C and 53% while on September 5 they were 22.6 C and 77%. See Table S1 of the SI for more details.

**Fig. 3.**
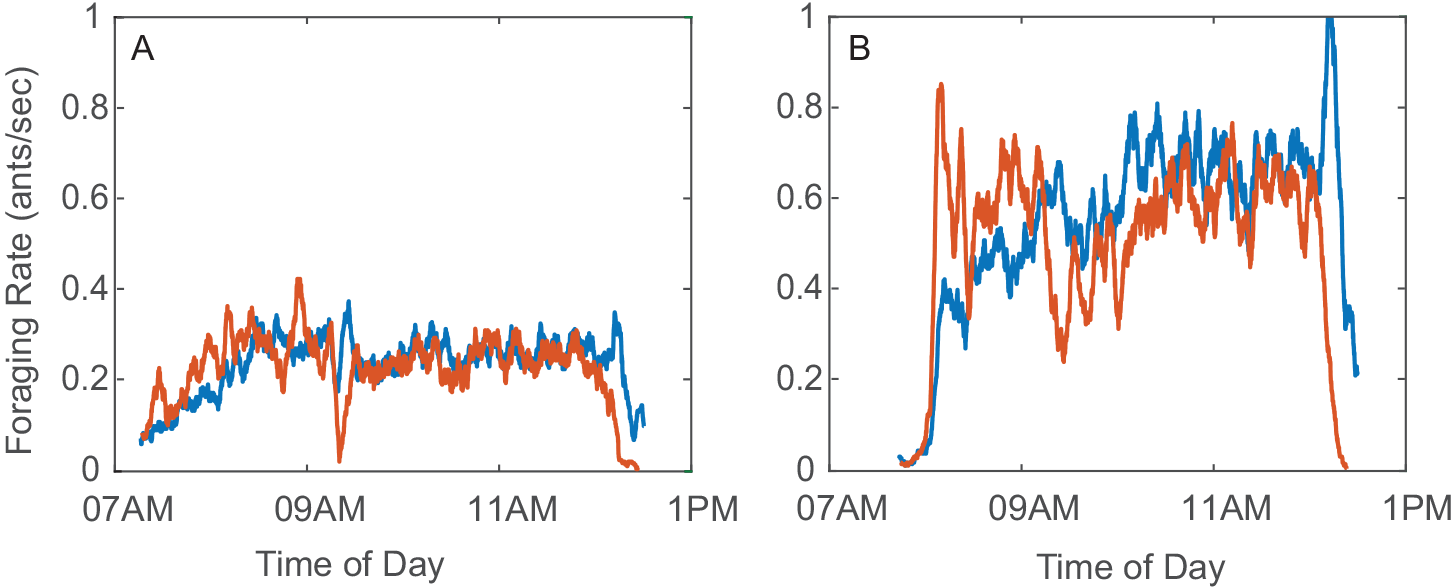
Plots of incoming foraging rate *r*_*in*_ (blue) and outgoing foraging rate *r*_*out*_ (red) versus time of day on August 20, 2016 for A) Colony 1357 and B) Colony 1317. The quasi steady-state (QSS) where incoming and outgoing rates equilibrate to a near-equal value can be observed for both colonies. The QSS rate for Colony 1317 is more than twice as great as it is for Colony 1357.

Fig. 3 and Fig. 4 are representative of observations that suggest the equilibration of incoming and outgoing foraging rates to a near-equal rate to be an important feature in the regulation of foraging in red harvester ant colonies. Further, the equilibrated rate, and the possibility of early cessation of foraging, depend on factors that differ among colonies (Fig. 3) and from day to day (Fig. 4). We examine the transient in foraging rates further in Fig. 4. Early in the foraging day, both *r*_*in*_ and *r*_*out*_ increase rapidly with *r*_*out*_ increasing more rapidly than *r*_*in*_. This leads to a rapid increase in the number of active foragers outside the nest. The rapid increase in both rates is followed by a decrease in *r*_*out*_ to the equilibrated near-equal value of the QSS (Fig. 4A and C) or to an early return of the ants to the nest (Fig. 4E).

Input-output plots show the relation between incoming and outgoing foraging rates Fig. 4B, D, and F. These figures show the same data as Fig. 4A, C, and E, respectively, but plot *r*_*out*_(*t*) versus *r*_*in*_(*t*) with time of day *t* in hours indicated by the color scale. The transient in rates during the early part of the foraging day appear as curved trajectories above the diagonal, because *r*_*out*_(*t*) is typically higher than *r*_*in*_(*t*). In Fig. 4B and D, the curve rises and then falls to the QSS value where the trajectory then equilibrates around a point on the diagonal corresponding to equal incoming and outgoing rates. This rise and fall of the curve in the input-output plot is typical, even when the trajectory returns to the origin as in the case of Fig. 4F.

The data shown in Figs 3 and 4 as well as in Fig. S2 are representative of the data collected in 2015, 2016, and 2017. Temperature and humidity for these data sets are given in Table S1 of the SI. Fig. S2B shows another example of a very early cessation of foraging. Fig. S2C and D show two different examples of long transients. Fig. S2E and F show two examples of a burst in the outgoing foraging rate at the start of the foraging day. See SI Appendix 2 for details.

**Fig. 4.**
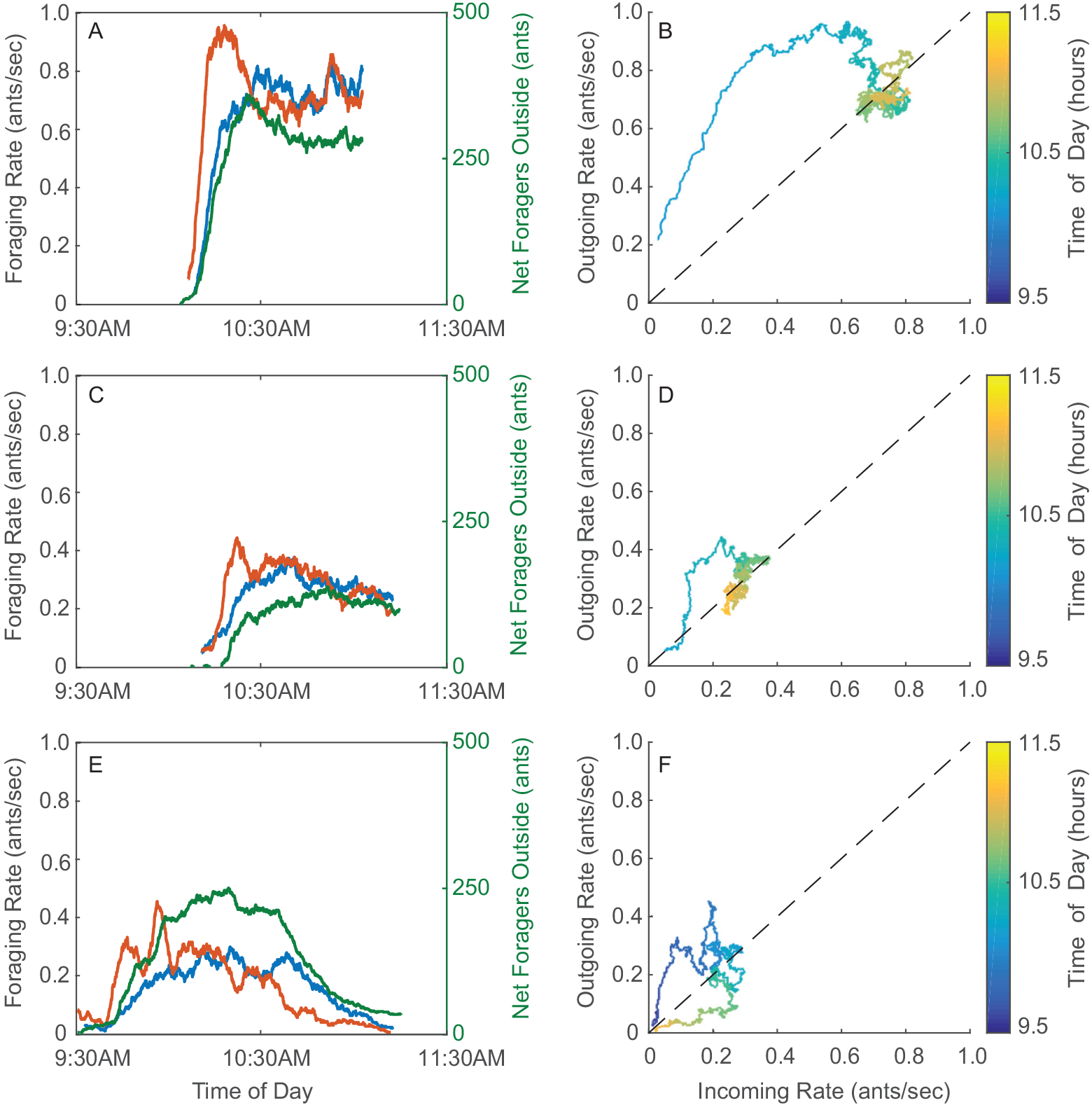
Plots of foraging rate data. Time series plots show incoming foraging rate *r*_*in*_ (blue), outgoing foraging rate *r*_*out*_ (red), and difference between the number of incoming and outgoing foragers (green) versus time of day. Input-output plots show *r*_*out*_(*t*) versus *r*_*in*_(*t*) with the color scale representing time of day *t*. A) and B) Colony 664 on August 27, 2015. C) and D) Colony 664 on August 31, 2015. E) and F) Colony 863 on September 1, 2015.

### Model Dynamics

#### Foraging Dynamics Inside the Nest

Given a sequence of incoming ants λ_*in*_, our open-loop model of foraging dynamics inside the nest (Fig. 1) predicts a corresponding sequence of outgoing ants λ_*out*_. We find an analytic approximation for the mapping from mean incoming foraging rate 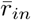 to mean outgoing foraging rate 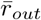, parametrized by volatility *c*. To do so, we assume λ_*in*_ is a Poisson process with (constant) mean incoming rate 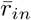; this is justified for observations of incoming and outgoing sequences of foragers for short periods of time [15].

We assign model parameter values to be *k* = 0.3, τ = 0.41, *ϵ*_1_ = 0.2, and *ϵ*_2_ = 0.05, which allow for rich dynamical behavior. While the qualitative behavior is unchanged for different values of *ϵ*_2_ ≪ 1, very high or low values of *k* and/or τ yield dynamics in which the stimulus *s* is either too low or too high to produce oscillations. So the values for *k* and τ are selected to balance their opposing effects on *s*.

For very low 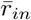, 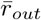 is low because *s* is low and the FN system remains in the resting state with occasional short-lasting periods of oscillatory behavior (Fig. S3). For very high 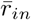, 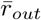 is also low because *s* is high and the FN system remains most of the time in the saturated state. In contrast, 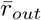 is high for 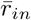 that yields an *s* that keeps the FN system inside the oscillating region. In the oscillating region, 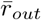 is equal to the frequency of the oscillations, which is inversely proportional to the volatility *c* as we show in SI Appendix 1. The oscillating region is given by the range of *s* between the FN bifurcation points *b*_1_ and *b*_2_, computed as 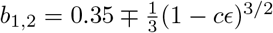. As the volatility *c* increases, the size of the oscillating region decreases (Fig S4).

To get an expression for the natural frequency of the oscillations in the FN, we compute an asymptotic approximation for its period *T*_*LC*_(*s*, *c*) as *ϵ*_2_ goes to zero, see SI Appendix 1 and Fig. S5. Under the assumption of a Poisson incoming rate, the process *s* is ergodic (see SI Appendix 3). Thus, over sufficiently long periods of time, suitable time statistics converge to ensemble statistics, allowing us to approximate the fraction of time that *s* spends in the oscillating region using 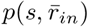, the probability density function of *s* at steady-state. We compute 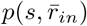 in SI Appendix 4 as a piecewise function where the piecewise elements satisfy recurrence equations and depend on *k* and τ. From this we can construct an analytical expression for 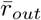 as a function of both 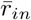 and *c* (see SI Appendix 3):

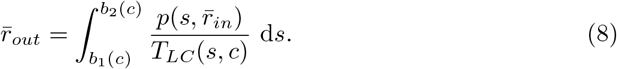

In Fig. 5A we plot 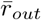 versus 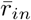 using Eq. (8) for different values of *c*. The resulting open-loop input-output curves, which we call *nest I/O curves* show that the analytic mapping from 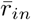 to 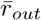 depends nonlinearly on *c*. The increasing steepness of the curve at low 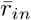 becomes more pronounced for higher *c* because the frequency of oscillations is proportional to *c*. Similarly, the decreasing steepness of the curves for high 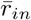 also becomes more pronounced for higher *c*. This is because as *c* increases *b*_2_ decreases, causing the FN to saturate at lower 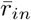 values. The maximum value of 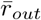 takes place at the 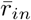 that yields an *s* that keeps the FN system inside the oscillating region. Because of this, the maximum 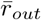 must be less than or equal to the natural frequency of the oscillations at the given value of *c*.

**Fig. 5.**
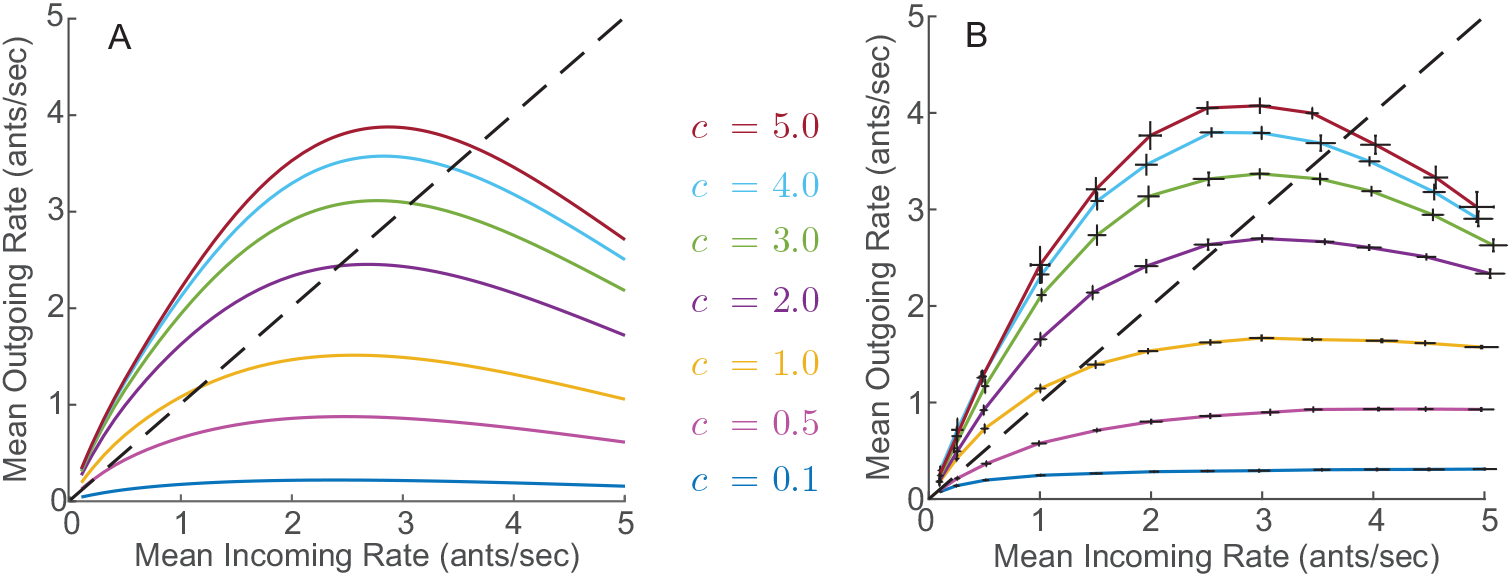
A) Analytical approximations for the nest I/O curves. B) Simulated nest I/O curves for different values of *c*. Each pair of error bars correspond to 10 simulation trials, each 5 minutes long, with a constant incoming rate 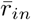 and constant volatility *c*. The dashed black line represents points at which 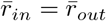.

In Fig. 5B we show the nest I/O curves obtained by simulating the open-loop system for different constant Poisson incoming rates at a fixed volatility. We measured the resulting mean outgoing rate in each case. We set λ_*in*_ to a five-minutes-long Poisson process, and, in each of 10 simulation trials, we recorded the output λ_*out*_. We computed the mean outgoing foraging rate 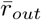 by dividing the total number of outgoing foragers in the trial by the 300 seconds that the trial lasted. We used the average of the 10 trials as a point estimate for 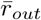 as a function of 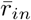 given the volatility parameter *c*. We constructed nest I/O curves by repeating this point estimation process for twelve different values of 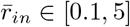 while keeping *c* constant.

The simulated I/O curves in Fig. 5B are in good agreement with the analytical I/O curves in Fig. 5A. The fact that there is good agreement between the simulation curves computed from short 5-minute-long input sequences and the analytical curves derived under the assumption of an infinite time period suggest that our analytical approximation is also valid across short timescales. We make use of this in our analysis of the closed-loop model dynamics.

The points at which the nest I/O curves in Fig. 5 intersect the black dashed diagonal line correspond to 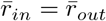, which are predictive of the (quasi) steady-state solutions at an equal incoming and outgoing foraging rate observed in the data. Fig. 5 suggests that for sufficiently high values of *c*, the equal foraging rate is positive and bounded away from zero, capturing a nontrivial steady-state foraging rate as in Fig. 3 and Fig. 4B and D. However, Fig. 5 suggests that for low values of *c*, the equal foraging rate is nearly zero, capturing a steady-state with negligible foraging as in Fig. 4F.

To understand first consider that, because *k* > *b*_1_, every incoming forager elicits at least one oscillation in the FN output and so at low 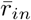, 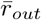 is equal to or larger than 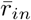. At high *c* values, the frequency of oscillations in the FN is high and a single incoming forager will elicit more than one oscillation, resulting in nest I/O curves with an initial slope higher than one and an intersection with the diagonal line at a single point away from the origin. In contrast, for low *c* values, a single incoming forager will elicit exactly one oscillation, resulting in nest I/O curves with an initial slope of one, i.e., the curve lies on the diagonal line close to the origin and intersects nowhere else.

We can approximate the value of *c* above which a single incoming forager results in more than one outgoing forager, guaranteeing an isolated nontrivial intersection of the nest I/O curve with the diagonal line. For *b*_1_ < *k* < *b*_2_, it can be shown that the number of oscillations caused by a single incoming forager is at most (–τ log *b*_1_/*k*)/*T*_*LC*_. We numerically solved this expression for *c* using the asymptotic expansion of *T*_*LC*_described in SI Appendix 1 and found that for *c* > *c*\* = 0.5287 the FN oscillates at least two times per every incoming forager. Therefore, *c* > *c*\* is a sufficient condition for the nest I/O curve to lie above the diagonal line at low 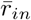 and to intersect the diagonal line at an isolated point, corresponding to a nontrivial steady-state foraging rate.

#### Foraging Dynamics Outside the Nest

Given a sequence of outgoing foragers λ_*out*_ with rate *r*_*out*_, the foraging dynamics outside the nest predicts a corresponding delayed sequence of incoming foragers λ_*in*_ with rate *r*_*in*_. We use results from queueing theory to find analytic expressions relating *r*_*out*_ to *r*_*in*_ and the expected number of active foragers outside the nest.

To facilitate the analysis we assume that λ_*out*_ is a non-homogeneous Poisson process (i.e., a Poisson process with time-varying rate) [15]. Applying known results for queues with a non-homogeneous Poisson distribution of arrival times [38] we obtain the following three results:

1. Let *Q*(*t*) represent the number of active foragers outside the nest, then, for each time *t*′ = *t*/60 where *t* is seconds, *Q*(*t*′) has a Poisson distribution with mean

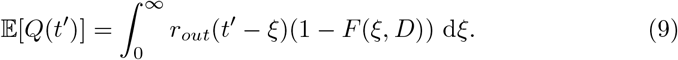
2. The output process describing how foragers leave the queueing system, that is, the process λ_*in*_ describing how foragers return to the nest, is a non-homogeneous Poisson process with mean

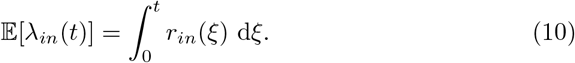
3. *r*_*in*_ is related to *r*_*out*_ by

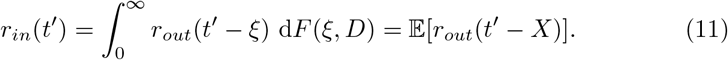

Eqs. (7) and (9) show how the active number of foragers outside the nest depends on the history of outgoing foragers. Eq. (10) shows that if the outgoing foraging process is a non-homogeneous Poisson process, then the incoming foraging process is also a non-homogeneous Poisson process. And Eq. (11) shows how the incoming foraging rate *r*_*in*_ depends on the history of the outgoing foraging rate *r*_*out*_.

At steady-state, the outgoing foraging rate is constant, i.e., *r*_*out*_(*t*) = *r*\*, and Eq. (11) reduces to *r*_*in*_ = *r*_*out*_ = *r*\*, i.e., the incoming foraging rate is also constant and equal to the outgoing foraging rate. Moreover, Eq. (9) reduces to

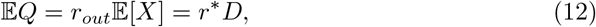

i.e., the mean number of active foragers outside the nest is given by the steady-state foraging rate *r*∗ multiplied by the average foraging trip time *D*.

The relaxation time for the queue output process to reach steady-state can be analyzed by considering the step-function arrival rate *r*_*out*_(*t*′) = 0 for *t*′ < 0 and *r*_*out*_(*t*′) = *r*\* for *t*′ ≥ 0. Then, from Eq. (11), 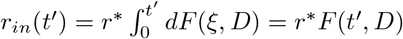 for *t*′ ≥ 0. The difference between the queue input and output rates as a function of time is

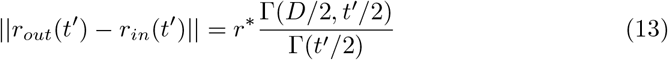

for *t*′ ≥ 0. For *D* = 2, the right-hand side of Eq. (13) simplifies to *r*\**e*^−*t*′/2^ and the foraging queue converges exponentially in time towards a steady-state where the input and output rates are equal.

#### Closed-loop Model Dynamics

In our model, outgoing foragers from the nest go out to forage, return to the nest as incoming foragers after finding a seed, and then go back out to forage again if sufficiently excited (Fig. 1). Here we show that adding the feedback connection from outgoing to incoming foragers to the open-loop dynamics in the nest yields long-term dynamics with a stable and attracting fixed-point where the incoming and outgoing rates are equal and robust to small perturbations in rates. When the volatility *c* is above *c*\*, the critical value studied above, the steady-state foraging rate is nontrivial, whereas if *c* is well below *c*\*, the steady-state foraging rate is negligible.

For the dynamics inside the nest, we have shown that *c* parametrizes a family of nest I/O curves, described by Eq. (8), which map 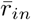 to 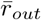. For *c* ≪ *c*\*, the nest I/O curve always has slope less than or equal to 1, such that it lies on or below the diagonal line where *r*_*in*_ = *r*_*out*_. For *c* > *c*\*, the nest I/O curve has initial slope greater than 1 and then intersects the diagonal line *r*_*in*_ = *r*_*out*_ at a nontrivial point. For the dynamics outside the nest, we have shown that the mapping from *r*_*out*_(*t*) to *r*_*in*_ (*t*) is described by a time delay given by Eq. (11).

We study the closed-loop model dynamics for timescales ranging from tens of minutes to hours by investigating the behavior of a discrete iterated mapping *r*_*n*_ = *G*_*c*_(*r*_*n*–1_) where *r*_*n*_ represents the mean foraging rate at time step *n*, and where the time in between steps is given by the time delay from *r*_*out*_(*t*) to *r*_*in*_(*t*) introduced by the foraging outside the nest. The mapping *G*_*c*_: ℝ_≥0_ → ℝ_≥0_ is defined by the c-dependent nest I/O curves shown in Fig. 5. *G*_*c*_ describes the process by which the incoming foraging rate becomes the outgoing foraging rate through the dynamics of forager activation inside the nest, which then becomes the incoming foraging rate at a later time.

When *G*_*c*_ lies below (above) the diagonal line where *r*_*in*_ = *r*_*out*_, the average number of outgoing foragers per every incoming forager is less (greater) than one, and iterations of *G*_*c*_ decrease (increase) *r* (Fig. 6). For *c* > *c*\*, *G*_*c*_ has one unstable fixed point at the origin and one attractive stable fixed point where *r*_*in*_ = *r*_*out*_. For *c* ≪ *c*\*, *G*_*c*_ has a small interval of fixed points close to the origin. Thus, the closed-loop model dynamics evolve in time towards either a finite steady-state foraging rate *r*_*in*_ = *r*_*out*_ = *r*\* (Fig. 6, *c* =2 and *c* = 5) or to negligible foraging (Fig. 6, *c* = 0.1).

**Fig. 6.**
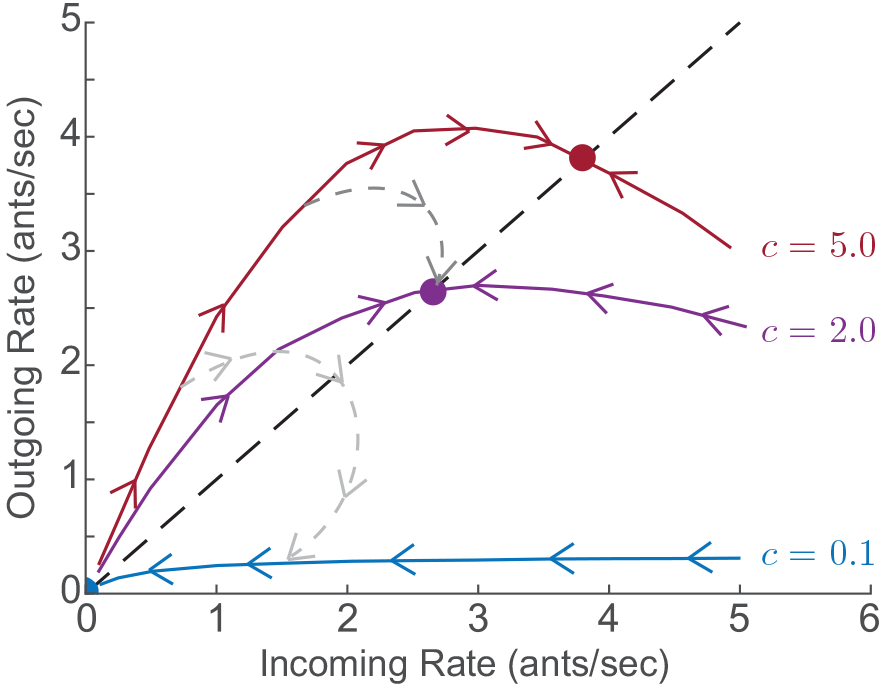
Model dynamics illustrating response of foraging rates to environmental conditions. Red, purple, and blue curves show closed-loop trajectories of *r*_*out*_(*t*) versus *r*_*in*_(*t*) for fixed volatility c equal to 5.0, 2.0, and 0.1, respectively. Initially, all available foragers are uninformed about the environment and have volatility *c*_*u*_ = 5.0. The darker gray dashed curve shows the dynamics in the case when foragers exposed to the environment reduce their volatility to *c*_*i*_ = 2.0, as might happen on a moderately hot and dry day. The lighter gray dashed curve shows the dynamics in the case when foragers exposed to the environment reduce their volatility to *c*_*i*_ = 0.1, as might happen on a very hot and dry day.

The robustness of the steady-state rates to small perturbations in λ_*in*_ results from the balance between positive feedback from incoming ants activating a larger number of outgoing ants, and negative feedback from saturation effects. The magnitude of the steady-state and the variance around it are positively correlated to the mean and variance of *s*, both of which increase with *c* for fixed *r*_*in*_ (see SI Appendix 3). The magnitude depends on *c*, *k*, and τ, given by Eq. (8), which can be numerically solved to find how the QSS foraging rate changes with *c* (Fig. 7A).

**Fig. 7.**
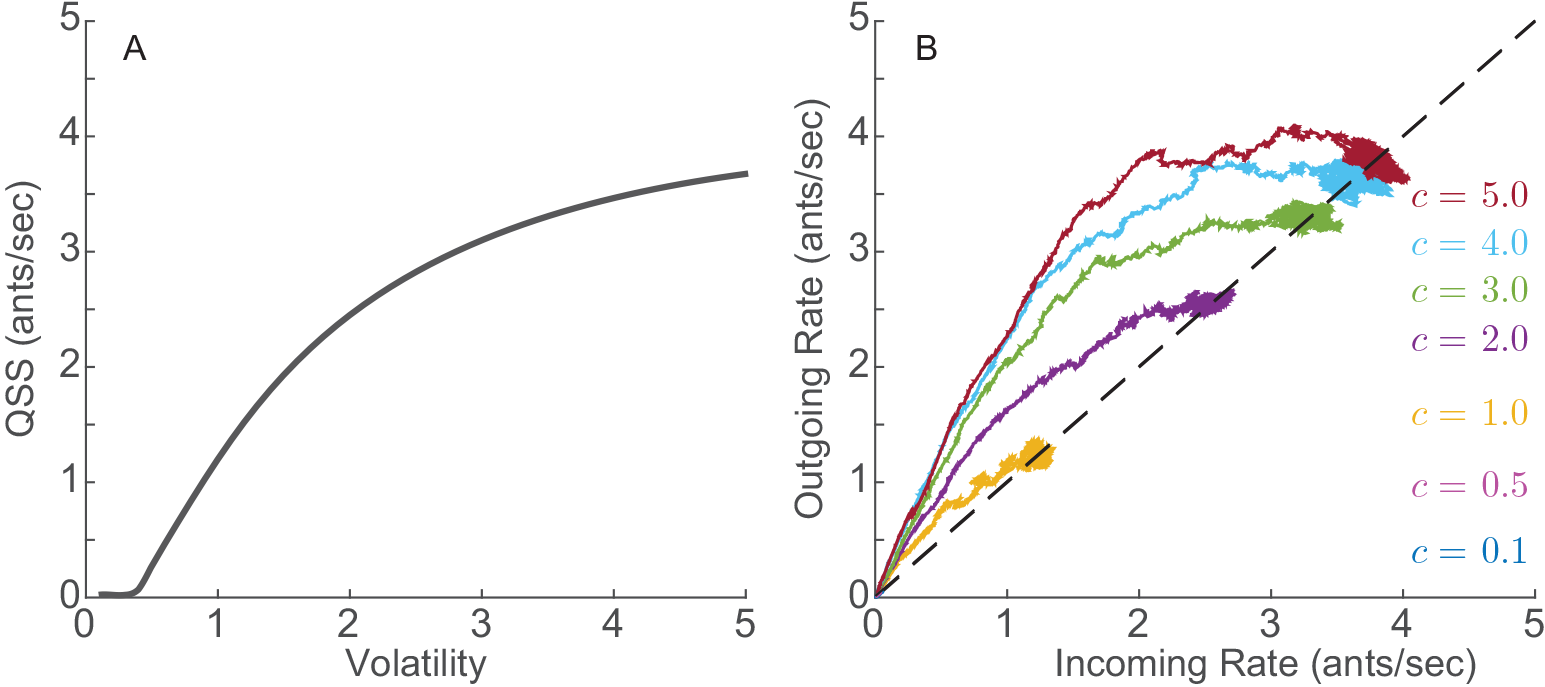
A. Analytical magnitude of the quasi steady-state (QSS) foraging rate obtained from numerically solving Eq. (8). B. Closed-loop model simulations for 7 different values of volatility *c*. The initial sequence of incoming foragers for all simulations was set equal to the first 11 minutes of the initial sequence of incoming foragers from Colony 859 on August 20, 2017 which has a mean incoming rate of 0.01 ants/sec. The total time for all simulations was 3 hours. The mean foraging time was set to 10 minutes (*D* = 10).

As shown in Fig. 7B, simulations of the closed-loop model validate the predictions of the iterated mapping model. We initialize the foraging dynamics by setting λ_*in*_ from *t* = 0 to *t* = 60(*D* + 1) to be equal to the first *D* + 1 minutes of the initial sequence of incoming foragers for Colony 859 on August 20, 2017, which has the very low mean incoming rate of 0.01 ants/sec during the first 15 minutes (panel C, Fig. S2).

#### Closed-loop Dynamics with Response to Environmental Conditions

For Poisson sequences of incoming foragers, the mean outgoing foraging rate of the colony is given as the weighted sum of the outputs of the uninformed and informed:

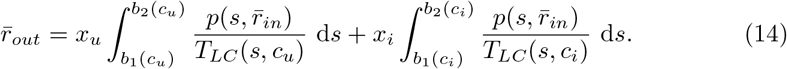

The closed-loop dynamics can still be studied as an iterated mapping, but we allow the mapping to evolve in time, *G*_*c*_ = *G*_*c*_(*t*), from an initial mapping 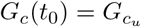 with volatility *c*_*u*_ to a final mapping 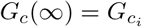 with volatility *c*_*i*_. We illustrate in Fig. 6) for *c*_*u*_ = 5.0 and *c*_*i*_ = 2.0 and for *c*_*u*_ = 5.0 and *c*_*i*_ = 0.1. The dynamics first evolve along 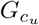 (red), but as *x*_*i*_ increases, the dynamics shift increasingly to 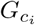, and the trajectory on the plot of *r*_*out*_(*t*) versus *r*_*in*_(*t*) moves towards the *c*_*i*_ curve. In the case *c*_*i*_ = 2.0, the trajectory converges to the fixed point associated with *c* = 2.0 (darker gray dashed curve). In the case *c*_*i*_ = 0.1, the trajectory converges to the only fixed point of 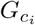, which is the origin, leading to a cessation of foraging (lighter gray dashed curve).

Fig. 8 shows the resulting time-series and input-output plots for three different simulations of the model with the mechanism for response to environmental conditions. The simulations are distinguished by the set of four parameters: *c*_*u*_, *c*_i_, *N*, and *D*. The simulated trajectories qualitatively resemble the trajectories from the field observations shown in Fig. 4. We set the initial sequence of incoming foragers as in Fig. 7B.

Fig. 8A and B show the results for *c*_*u*_ =3, *c*_*i*_ = 0.9, *N* = 500, and *D* = 5. In this case, *c*_*u*_ is much higher than *c*_*i*_, leading to a system with an overshoot behavior in which the outgoing foraging rate increases more rapidly than the incoming rate and then decreases before settling around a steady-state where the rates are approximately equalto 0.7 ants/sec. This is qualitatively similar to the observations of Colony 664 on August 27, 2015 of Fig. 4A and B. The net number of foragers outside the nest at steady-state fluctuates with low variability at around 230, close to the prediction given by Eq. (12).

**Fig. 8.**
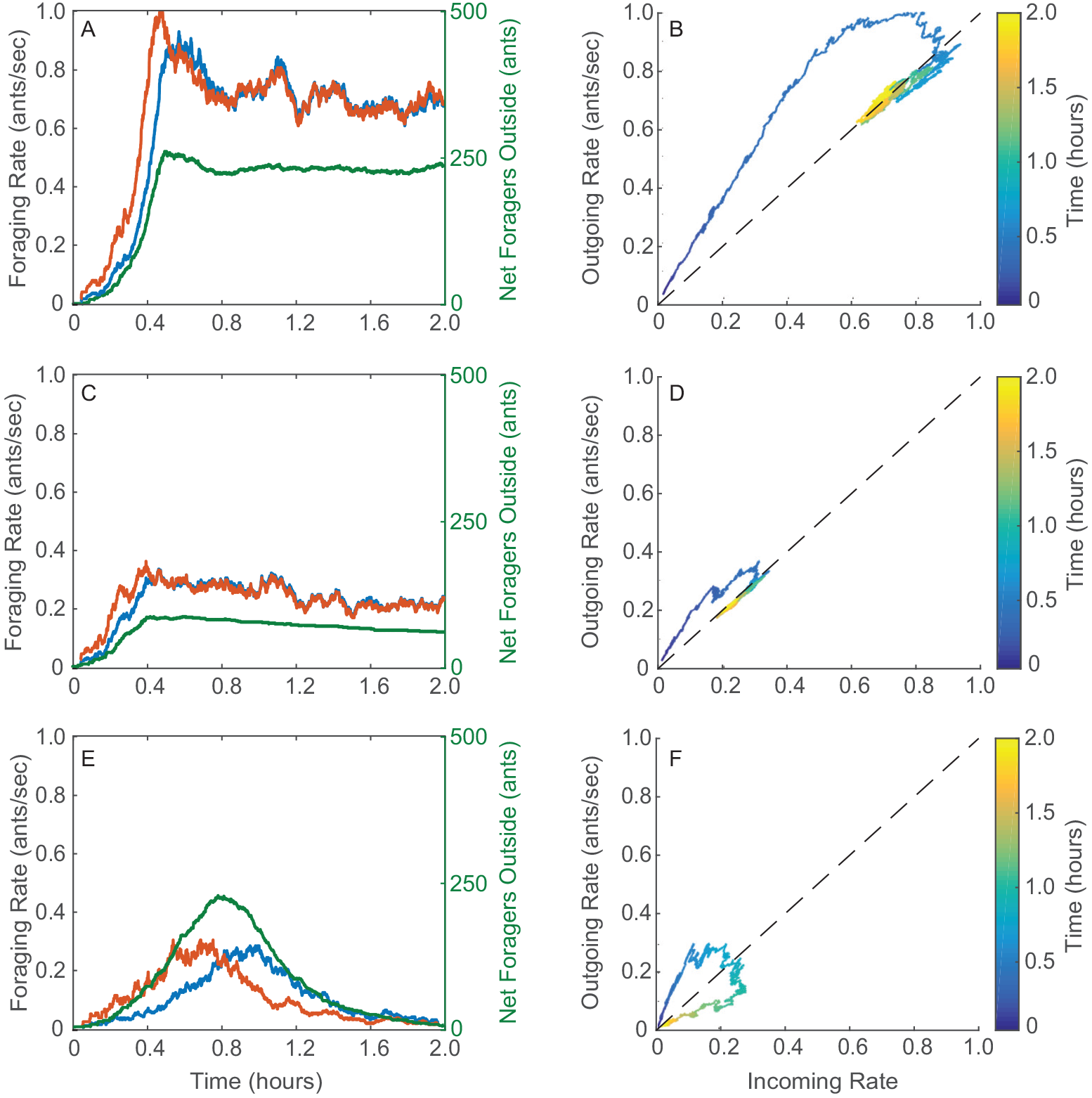
Simulations of the closed-loop model with the adaptation mechanism. Plots are of the same form as in Fig. 4, and qualitative comparisons can be made between A and B here and Fig. 4A and B, between C and D here and Fig. 4C and D, and between E and F here and Fig. 4E and F. A) and B) *c*_*u*_ = 3, *c*_*i*_ = 0.9, *N* = 500, *D* = 5. C) and D) *c*_*u*_ = 3, *c*_*i*_ = 0.75, *N* = 200, *D* = 5. E) and F) *c*_*u*_ = 5, *c*_*i*_ = 0.02, *N* = 600, *D* = 15.

Fig. 8C and D show the results for *c*_*u*_ =3, *c*_*i*_ = 0.75, *N* = 200, and *D* = 5. This case simulates the same colony as in Fig. 8A and B but on a hotter and drier day, when the total number of ants *N* that engage in foraging may be reduced and the volatility of the informed ants *c*_*i*_ may be reduced. The overshoot behavior is followed by the foraging rates settling around a steady-state of about 0.25 ants/sec. This is qualitatively similar to the observations of Colony 664 on August 31, 2015 of Fig. 4C and D.

Fig. 8E and F show the simulation results for *c*_*u*_ = 5, *c*_*i*_ = 0.02, *N* = 600, *D* = 15. In this case, *c*_*i*_ is close to zero, leading to a colony that goes out to forage but then returns to the nest without sustained foraging. This is qualitatively similar to the observations of Colony 664 on August 27, 2015 of Fig. 4E and F.

The time it takes for the colony to transition from fully uninformed to fully informed is dictated by *c*_*u*_, *c*_*i*_, *D*, *N*, and the initial condition for *r*_*in*_ and *r*_*out*_. For low values of *c*_*u*_, the outgoing foraging rate at the start of the day is low and the corresponding rate at which foragers become informed is low too. If *c*_*i*_ is high, the colony will spend a long time at a low rate before the number of informed foragers is sufficiently high to increase the foraging rates. We show a simulation of high *c*_*i*_ in Fig. S6A, which can be compared with the data from Colony 859 on August 20, 2017, plotted in Fig. S2C.

Low values of *c*_*i*_ can cause long transients. Once a critical number of foragers has become informed, it will be difficult for the remaining foragers to become informed since the colony behaves as an informed colony with low volatility. High *D* and *N* can also result in long transients because the time it takes for the colony to transition into a fully informed state depends on how many available foragers there are and on how long it takes for informed foragers to return to the nest. High *D* increases the capacity of the foraging queue, requiring higher numbers of active foragers at steady-state (see Eq. (12)). We show a simulation of high *D* and *N* in Fig. S6B, which can be compared with the data from Colony 1107 on August 16, 2017, plotted in Fig. S2D.

The initial conditions also affect the length of the transient. For an initially high value of *r*_*out*_, the number of active foragers increases very rapidly, reducing the time it takes for the colony to reach the informed state with foraging rates that reach a QSS. We show a simulation for an initially high value of *r*_*out*_ in Fig. S6C, which can be compared to data from Colony 1017 on August 23, 2016 and Colony 1015 on August 18, 2016, plotted in Fig. S2E and F.

### Discussion

We have derived and analyzed a low dimensional analytical model of foraging dynamics that requires only a small number of parameters to qualitatively capture a wide range of transient and steady-state features observed in the foraging rates of red harvester ant colonies. The model is distinguished from previous work in that it accounts for how incoming and outgoing rates, to and from the nest, change over long timescales, from tens of minutes to hours. Importantly, the long timescales allow for a model-based investigation into how a colony, with no centralized control and little individual information about the state of the colony or environment, can regulate its foraging rates to be robust to small disturbances and responsive to temperature and humidity outside the nest across minutes to hour-long timescales. Further, because the model is analytically tractable, it can be used to systematically derive predictions of foraging behavior as a function of critical model parameters and to explain variation in foraging activity among colonies in response to changing conditions.

Our model and analysis highlight the importance of feedback in the regulation of foraging activity. Previous work isolates the open-loop dynamics inside the nest, which maps incoming ants to outgoing ants on very short timescales. We address the minutes to hour-long timescales by examining analytically the closed-loop dynamics that connect the foraging activity outside the nest to the activation of foragers inside the nest through feedback generated by the ants themselves and their interactions with others. The stream of foraging ants out of the nest is the input to the foraging activity, and the output of the foraging activity is the stream of foraging ants into the nest.

Using analytical tools from dynamics and control theory, we show how the incoming and outgoing foraging rates first undergo a transient in which the outgoing rate exceeds the incoming rate, how the incoming and outgoing rates then stably approach a common value that is lower than the peak outgoing rate, and how the rates remain equilibrated at a quasi steady-state, despite disturbances, until mid-day when foraging rates go to zero. We show further how the transient and quasi steady-state rates differ under different foraging conditions, and how sensitivities to a small number of parameters can explain differences among colonies in the regulation of foraging.

To examine the underlying mechanisms that help explain changes over minute to hour-long timescales and differences over colonies in foraging rates, we isolate the parameter c in our model, which we call “volatility”, and we explore its role in foraging dynamics. Volatility approximates the average sensitivity of available foragers in the nest to interactions with returning foragers: the higher the c the fewer interactions needed to activate available foragers to go out and forage. The relationship between c and the activation of foragers is nonlinear, and the subtleties of our model reflect some of the complexities of the system. We use analytical predictions to show how c determines three important features of the foraging model dynamics (see Fig.7): 1) the initial transient in incoming and outgoing foraging rates, parametrized by *c*, 2) the equilibration of incoming and outgoing foraging rates to a stable, and thus robust, quasi steady-state rate, parametrized by *c*, and 3) the prediction of an early cessation of foraging or no foraging at all if *c* ≪ *c*\*, a critical volatility value *c*\*.

The behavior of different colonies on the same day or the same colony on different days thus correspond in the model to different values of volatility *c*. Lower values of *c* are consistent with hotter and drier days, because lower *c* means that available foragers are less volatile and thus less likely to go out and forage. Higher values of *c* are consistent with cooler and more humid days, because higher *c* means that available foragers are more volatile and thus more likely to go out and forage.

Because the temperature and humidity inside the nest remain relatively constant over the foraging day (Fig. S1), it is likely that foragers learn about the current outside conditions only after they have been on a foraging trip and thus exposed to the environment. To account for this in our model of a single colony on a single day, we implement two volatility terms *c*_*u*_ and *c*_*i*_ to distinguish between the volatility of “uninformed” available foragers in the nest who have yet to go on a foraging trip and the volatility of “informed” available foragers in the nest who have already been outside the nest on at least one foraging trip (Fig. 2). The result is a transition from the foraging activity of ants with volatility *c*_*u*_ to the foraging activity of ants with volatility *c*_*i*_, which can last from minutes to hours as each of the total *N* ants goes out at a different time on its first foraging trip and returns to the nest after foraging for an average of *D* minutes (Fig. 6).

We show that over a range of values for the four parameters *c*_*u*_, *c*_*i*_, *N*, and *D*, the model describes the range of transient and quasi steady-state foraging rate behaviors observed in the data collected for red harvester colonies in August and September of 2015, 2016, and 2017. In each of the data sets plotted in Fig. 3 and Fig. 4, the outgoing foraging rate initially increases faster than the incoming foraging rate and then the two rates equilibrate at a quasi-steady state. In Fig 4 B, D, and F, this behavior is described by outgoing-vs-incoming rate curves that initially rise above the diagonal line before converging onto the diagonal line where the rates are equal and lower than the maximum outgoing rate during the transient. That our model explains the shapes of these curves, and distinguishes the three different cases of Fig. 4 only through different values of the four parameters can be seen in the model simulations of Fig. 8. Fig. 4 B and D show the curves for foraging rate data corresponding to the same colony on two different days, the second day warmer and drier than the first. The simulations shown in Fig. 8 B and D recover the two differently shaped curves of Fig. 4 B and D, respectively, only by using different values of *c*_*i*_ and *N* that are consistent with the different environmental conditions. The data in Fig. 4 F show a colony discontinuing its foraging after the initial transient and the shape of this curve is recovered in the model simulation Fig. 8 F. Similar comparisons can be made for the data curves plotted in Fig. S2 and the model simulation curves plotted in Fig. S6.

The model represents the case in which foragers make an adjustment to their volatility only after their first foraging trip. To include more variability within a colony the model can be generalized to *M* > 2 groups of available foragers in the nest, distinguished by *M* values of volatility *c*_1_, …, *c*_*M*_. For example, the generalization can be used to account for foragers that make adjustments to how they respond to interactions in the nest after subsequent foraging trips due to repeated exposure or changing temperature and humidity. The generalization can also be used to account for decay of information for those foragers who stay in the nest for a long period after a foraging trip, or to represent foragers that return to the deeper nest after exposure to hot and dry outside conditions.

Our biologically informed, low-dimensional, and simply parameterized model allows for systematic exploration of mechanisms and sensitivities that can explain collective behavior and guide further theoretical and experimental investigations. Our use of well-studied excitability dynamics opens the way for comparison with other complex systems, such as neuronal networks, that are driven by excitable dynamics. The model together with our analysis based on dynamics and control theory contribute to a better understanding of the role of feedback across multiple timescales in the emergence of adaptive collective behavior in complex systems.

## Acknowledgments

We thank many dedicated undergraduate field assistants for their help: Sam Crow, Eleanor Glockner, Christopher Jackson, Sarah Jiang, Ga-Il Lee, Arthur Mestas, Becca Nelson, and especially Rebia Khan.

## Supporting Information

**Fig. S1.**
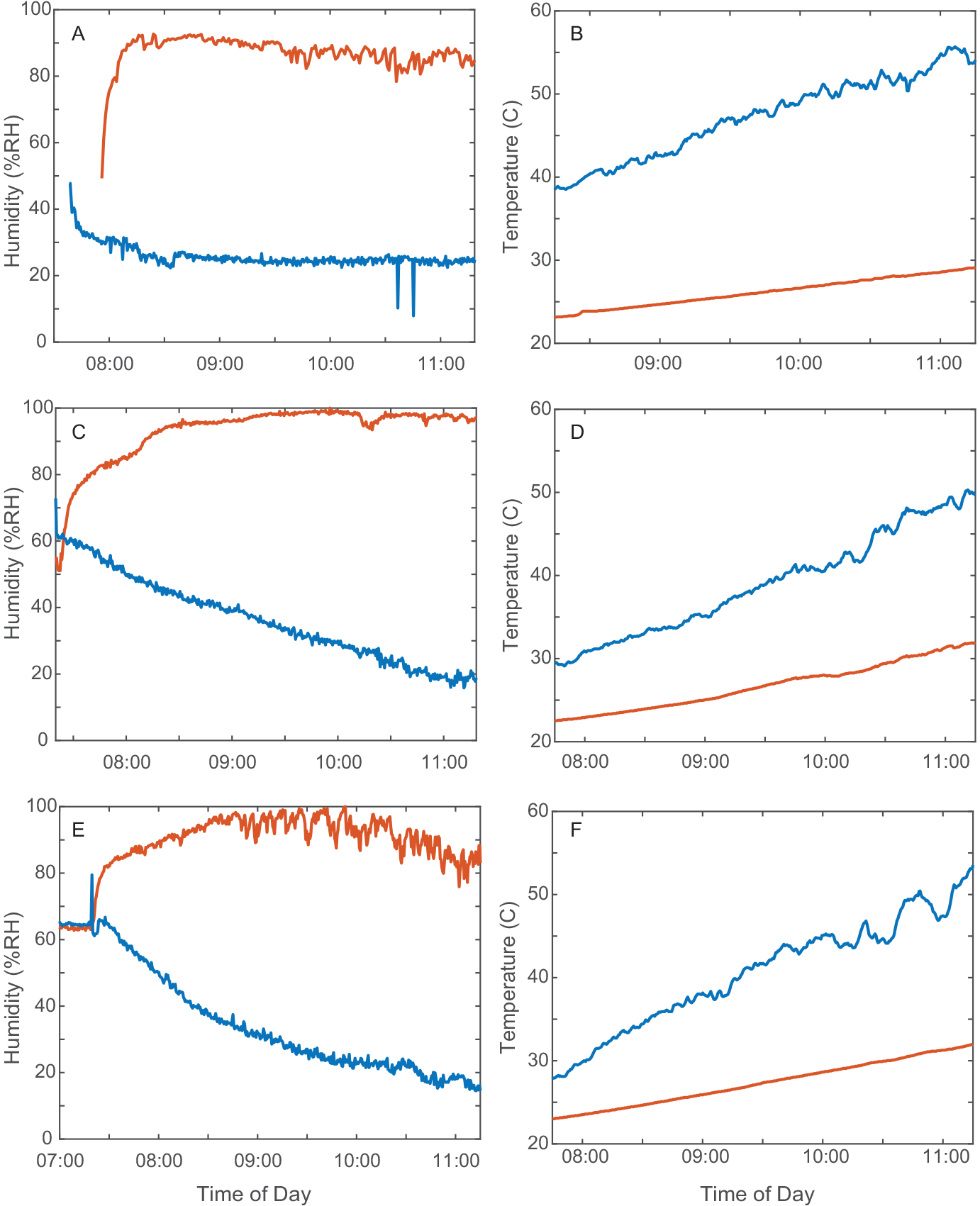
Humidity and temperature at surface of desert soil and inside the nest entrance chamber. Humidity and temperature readings recorded on the surface of the desert soil (blue) and inside the nest entrance chamber (red). Temperature and humidity ibutton sensors were placed outside but close to the nest entrance on the desert soil, unshaded, and inside in the nest in an excavated hole, which had been uncovered by excavation and then covered with glass on top and shaded. The humidity and temperature outside the nest change significantly throughout the morning hours while the humidity and temperature inside the nest entrance chamber remain relatively constant. The measured moderate rise in temperature inside the nest is likely due to the light coming into the nest entrance chamber through the glass. A) Humidity on August 29, 2014 (Colony E). B) Temperature on August 29, 2014 (Colony E). C) Humidity on August 31, 2015 (Colony 10). D) Temperature on August 31, 2015 (Colony 10). E) Humidity on September 1, 2015 (Colony 10). F) Temperature on September 1, 2015 (Colony 10).

**Fig. S2.**
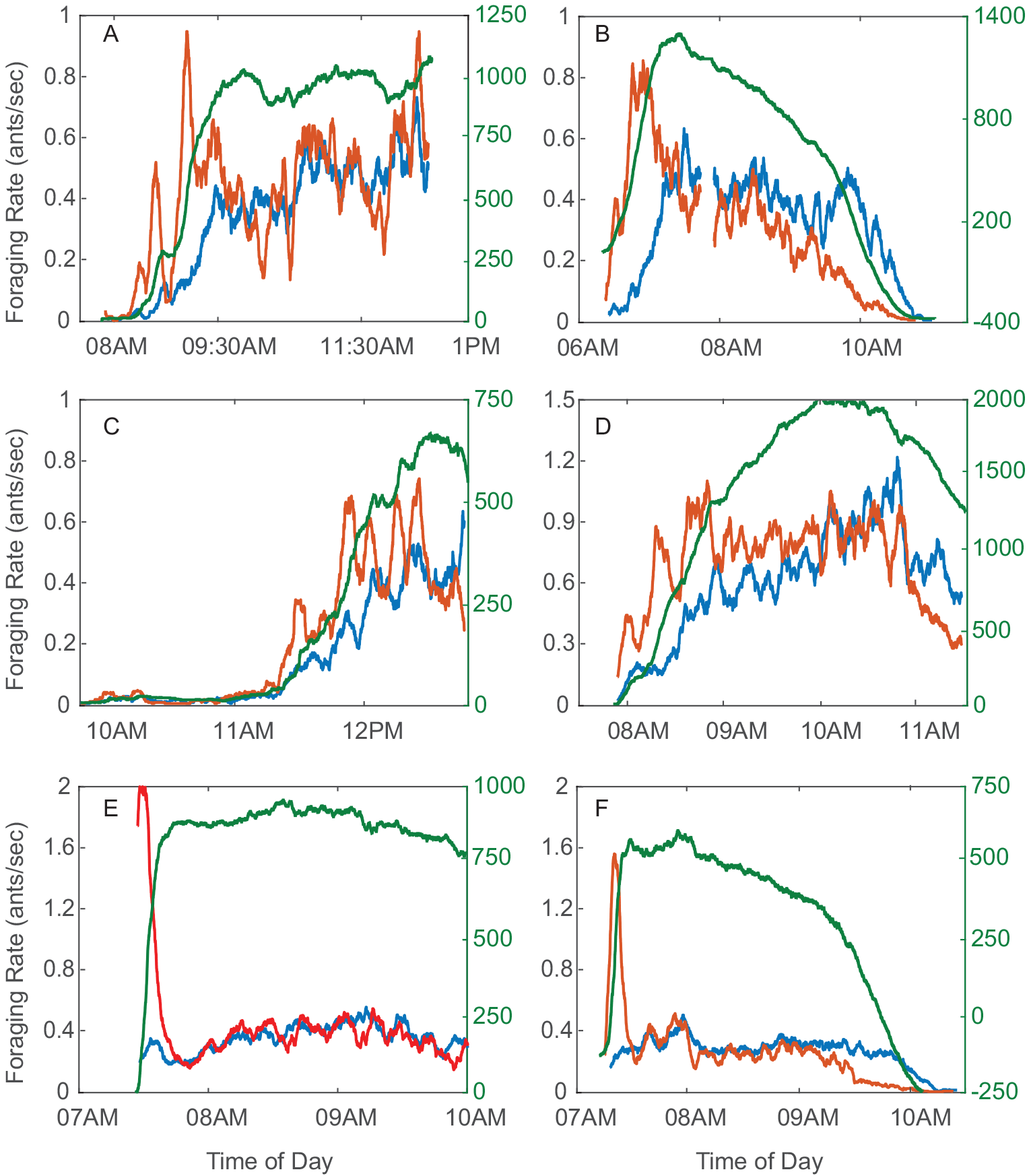
Additional field observations of foraging rates. Incoming foraging rate *r*_*in*_ (blue), outgoing foraging rate *r*_*out*_ (red), and difference between number of incoming and outgoing foragers (green) versus time of day. A) Colony 863 September 5, 2015 reaches a QSS at a high rate; compare to Fig. 4E when on the much hotter and drier day, September 1, 2015, Colony 863 returned to the nest early. B) Colony D19 August 08, 2016 returned to the nest early; the day was very hot and dry. C) Colony 859 August 20, 2017; the transient starts late in the morning. The day was cool and humid. D) Colony 1107 August 16, 2017; the transient is slow. The day was dry. E) Colony 1017 August 23, 2016; the initial transient is more like a burst of outgoing foragers. The day was dry. F) Colony 1015 August 18, 2016; another initial burst of outgoing foragers. The day was very dry.

**Table S1.**
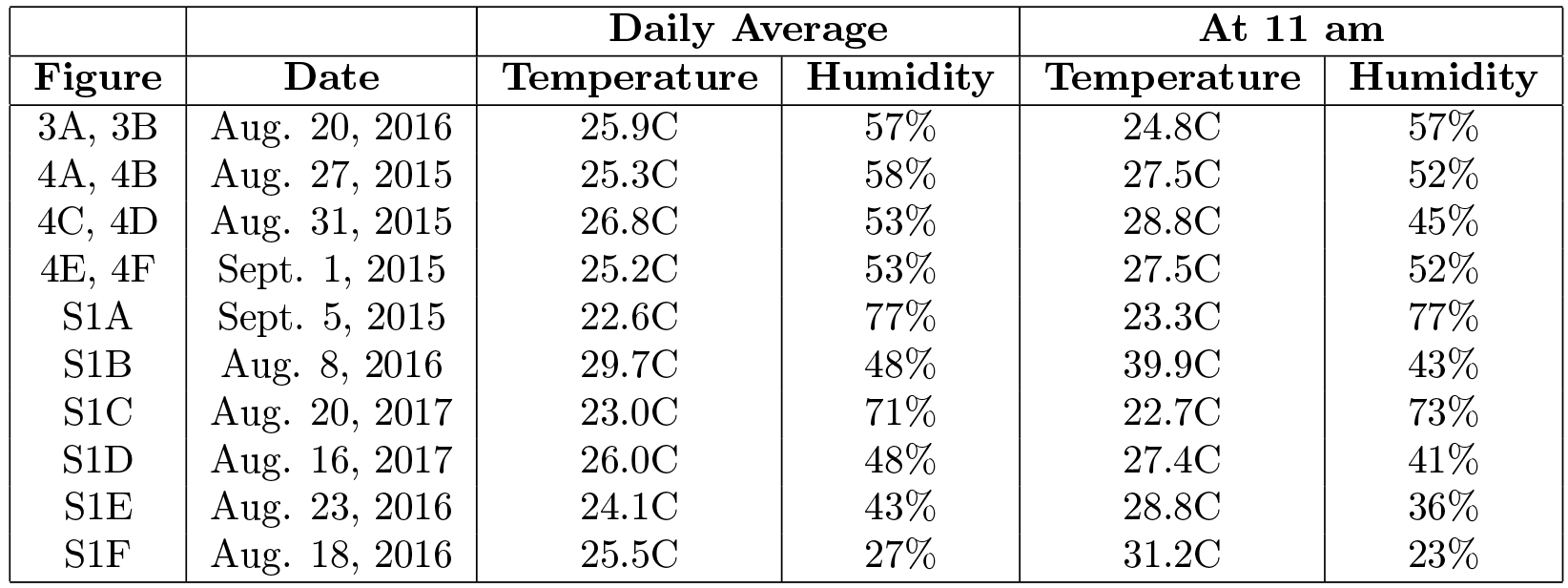
Temperature and relative humidity in Rodeo, New Mexico. Average temperature, average relative humidity, temperature at 11 am, and relative humidity at 11 am in Rodeo, New Mexico, USA for days with data plotted in Fig 3, Fig. 4, and Fig. S2. Data collected by the Citizen Weather Observer Program station E8703 and accessed through Weather Underground [39]. The station is located 1.7 miles from the study site.

**Fig. S3.**
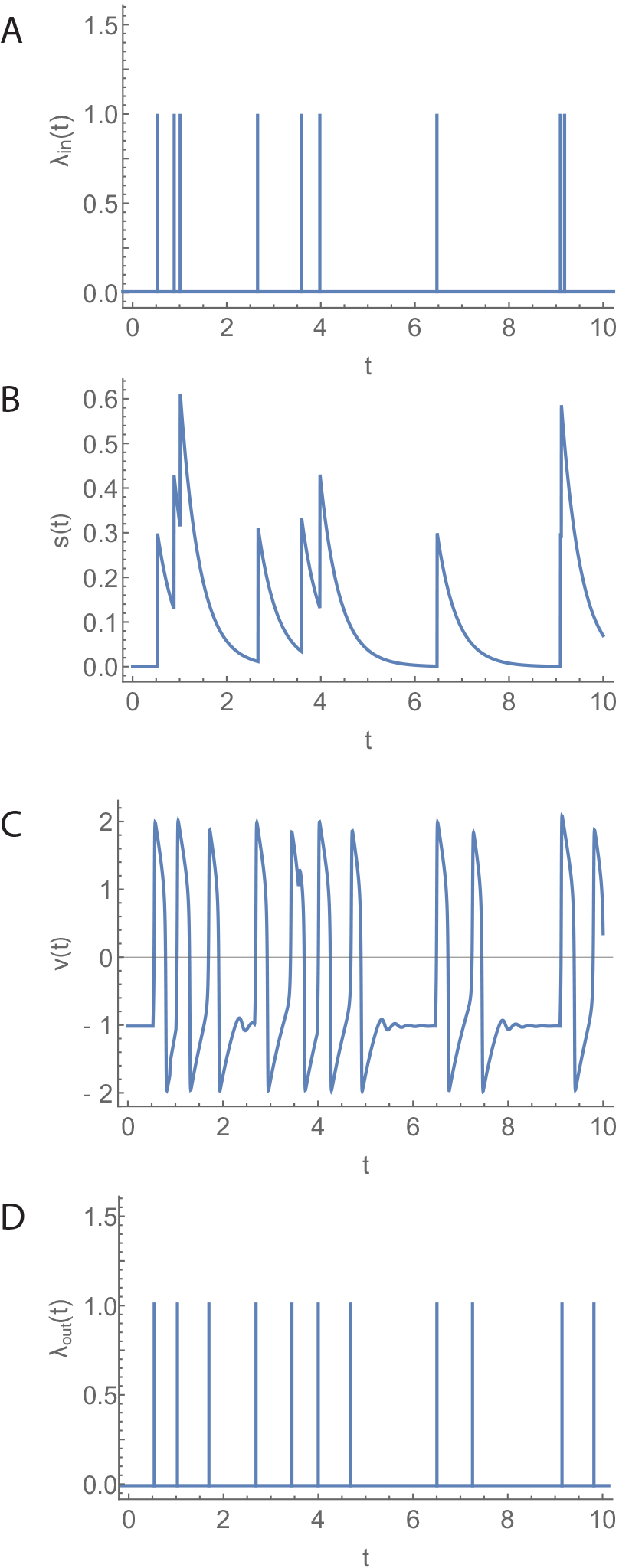
Open Loop Model. A) Poisson sequence of incoming foragers λ_*in*_. B) Stimulus signal *s* associated with λ_*in*_. C) FN output for input *s*. D) Sequence of outgoing foragers λ_*out*_ obtained by thresholding FN output from below at 0.75.

**Fig. S4.**
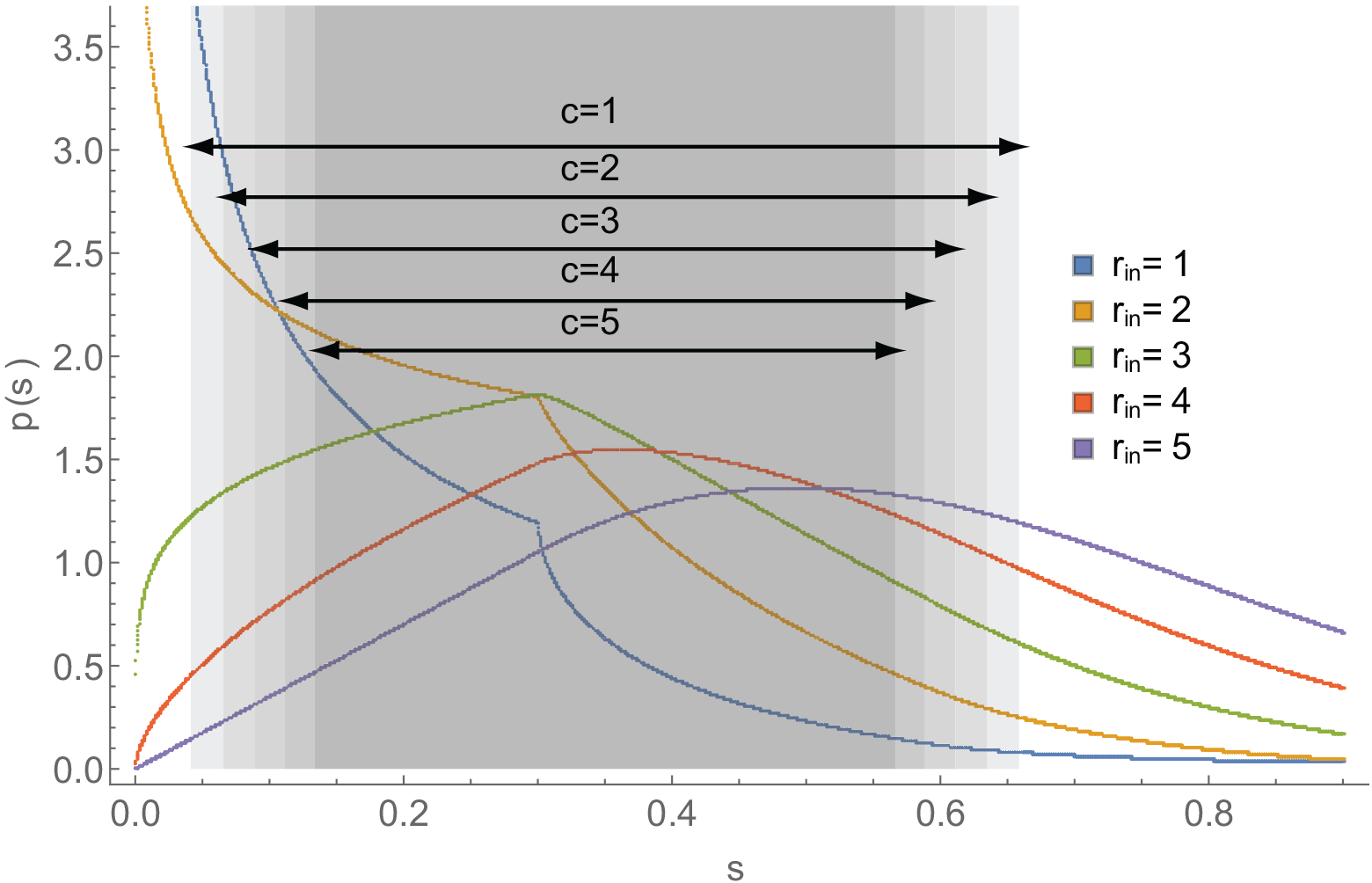
Probability density function for the stimulus function. Each curve represents the PDF *p* of the stimulus function *s* for different values of incoming rate *r*_*in*_. The gray rectangles represent the size of the oscillatory region in the F-N system (*b*_1_, *b*_2_) for different values of volatility *c*. For all curves, *k* = 0.3, τ = 0.41.

**Fig. S5.**
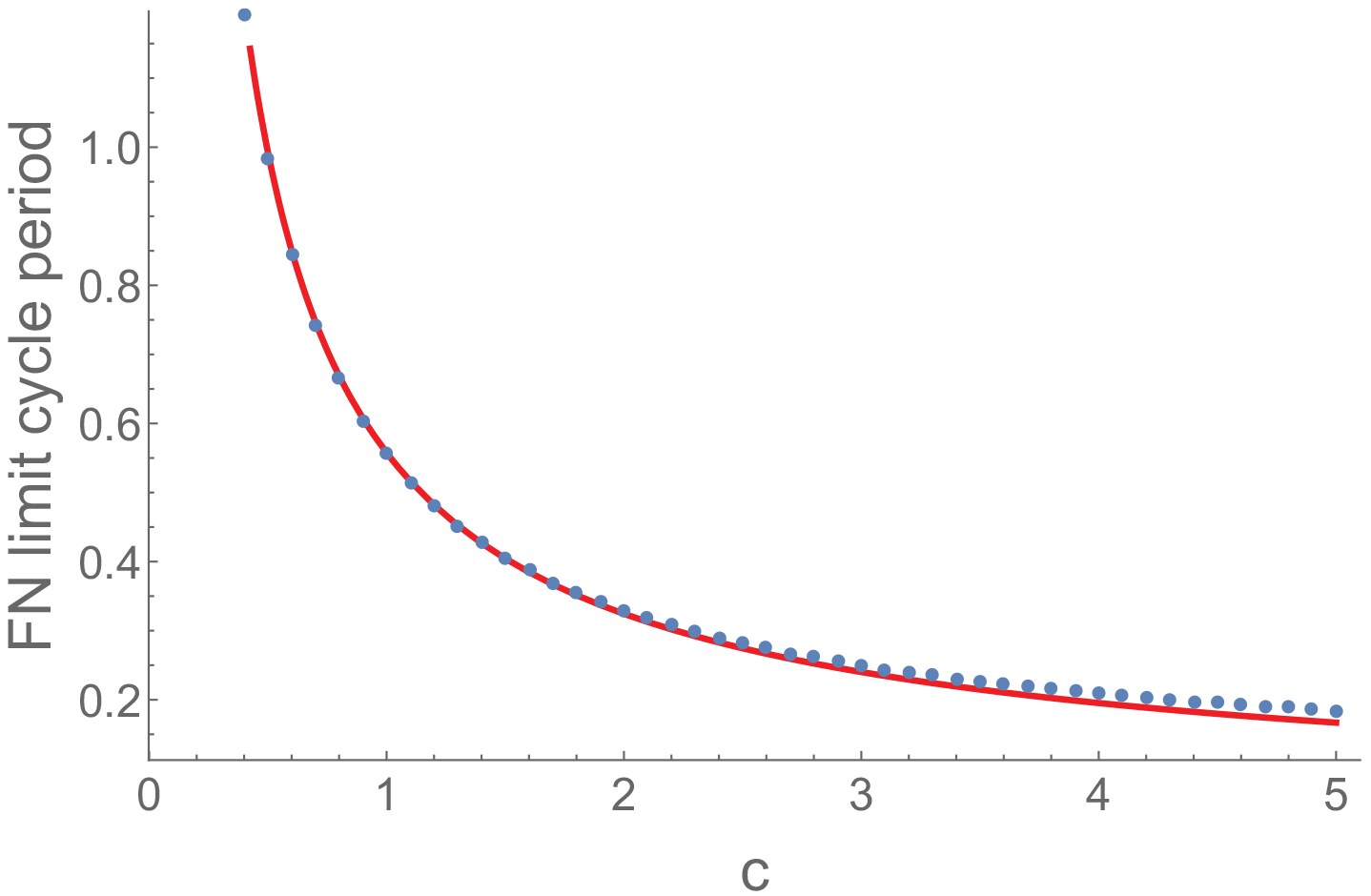
Period of FN Limit Cycle when s=0.35. Blue dots represent numerical simulations for the period of the FN limit cycle. The red curve represents the analytical approximation in SI Appendix 1. In both cases we set *s* = 0.35.

**Fig. S6.**
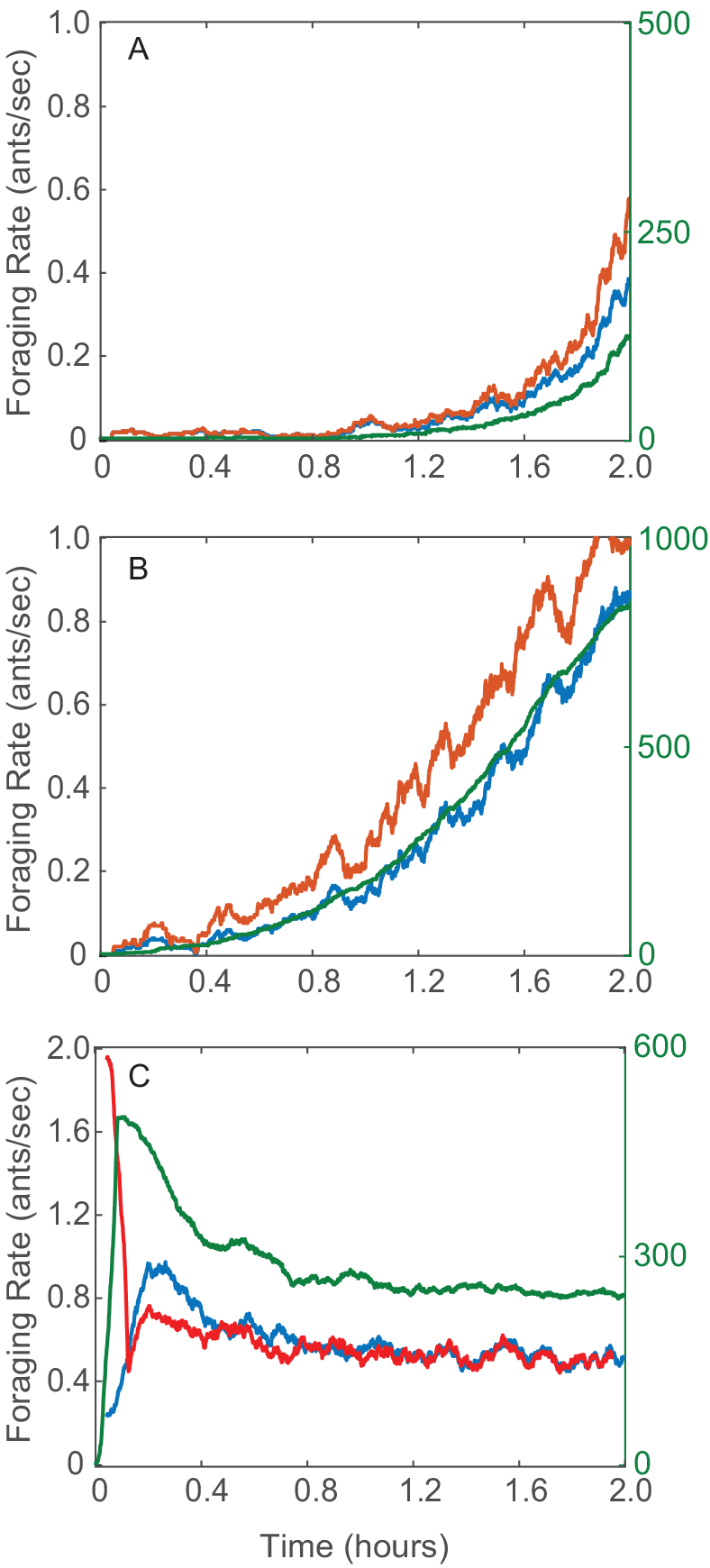
Additional simulations of the closed loop system with the adaptation mechanism. Plots resemble observed foraging behaviors in Fig. S2. Qualitative comparisons can be made between A here and Fig. S2C, between B here and Fig. S2D, and between C here and in Fig. S2E and F. A) *c*_*u*_ = 0.9, *c*_*i*_ = 2.2, *N* = 500, *D* = 5. Setting *c*_*u*_ < *c*_*i*_ where *c*_*u*_ is close to *c*\* results in a long period before the rates ramp up. B) *c*_*u*_ = 1, *c*_*i*_ = 1, *N* = 1000, *D* = 15. Setting the mean foraging trip time *D* to be large results in long lasting transients. C) *c*_*u*_ = 0.7, *c*_*i*_ = 0.9, *N* = 1000, *D* = 7. Setting the initial λ_*in*_ equal to sequence from the first 5 minutes of λ_*in*_ for Colony 1017 on Aug. 23, 2016 (see Fig. S2E) yields the behavior that follows an initial burst of foragers.

## SI Appendix 1

### Effect of volatility on the frequency of oscillations in the FN

Here we use results from [2] Chapter III, Theorem 3 to obtain an asymptotic expansion for the period of the limit cycle *T*_*LC*_ as ϵ_2_ goes to zero. We show that the period of the FN limit cycle is inversely proportional to *c* by computing the leading term in the expansion.

The limit cycle of the FN is comprised of four components: two fast components that stretch along the υ direction between the crest and valley of the cubic nullcline, and two slow components that stretch along the sides of the cubic nullcline. Because it takes much longer to traverse the slow components of the limit cycle than the fast components of the cycle, the period can be approximated by the time it takes trajectories to traverse the two slow components. These slow components are proportional to the length of the sides of the cubic nullcline, which we show are proportional to *c*.

#### Theorem 1.

*The limit cycle of the FN system*

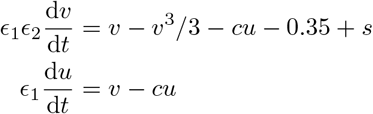

*has the asymptotic representation:*

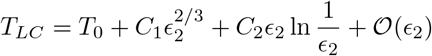

*as* ϵ_2_ → 0, *where C*_1_ *and C*_2_ *are constants and where*

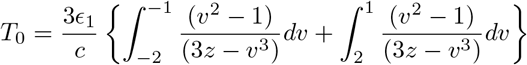

*Proof.* By Chapter III, Theorem 3 of [2], the limit cycle of the FN model has the asymptotic representation

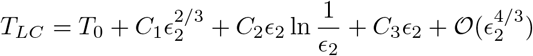

Or, equivalently,

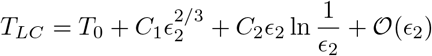

Let time be scaled by 1/ϵ_1_ and let *z* = *s* − 0.35, leading to the new system

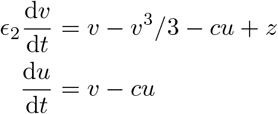

The critical manifold of this fast-slow system is *M*_0_:={(*υ*, *u*) ∈ ℝ^2^|*u* = (*υ* − *υ*^3^/3+*z*)/*c*}. In the limit ϵ_2_ → 0, the slow manifold is equal to the critical manifold. Let Ψ_0_ denote the limit cycle in this limit.

Using the description of *M*_0_ as a graph *u* = *h*(*υ*), the dynamics of the system on the slow flow can be written as

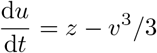

We get a second expression for d*u*/d*t* by differentiating *M*_0_ with respect to *t*

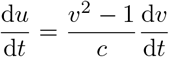

Equating the two expressions we obtain

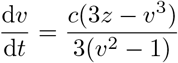

Multiplying both sides by d*t* and integrating over Ψ_0_,

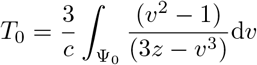

In Ψ_0_, the fast components of the orbit take place instantaneously and the time taken to complete the orbit is equal to the time taken to traverse the slow components. The slow components of the trajectory take place on the slow manifold between *υ* ϵ [−2, −1] and *υ* ϵ [1,2], yielding the expression

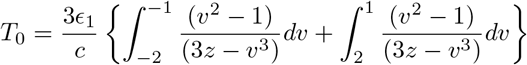

where time has been scaled back to its original form. This expression is inversely proportional to *c*. Furthermore, this integral has a short closed form solution when the slow components are symmetric (i.e. *z* = 0, or *s* = 0.35),

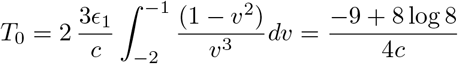

We compute the constants *C*_1_ and *C*_2_ by applying formulas 7.9 and 7.10 of [2] Chapter III, Theorem 3. When *s* ≠ 0.35, the flow along the system is not symmetric and the asymptotic representation is

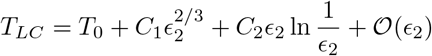

where

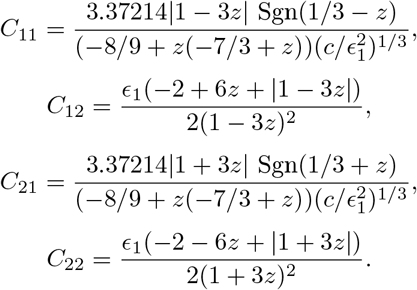

and Sgn represents the sign function.

When *s* = 0.35, *z* = 0 and the asymptotic representation becomes

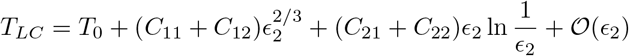

where

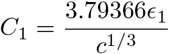

and

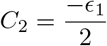

## SI Appendix 2

### Additional field observations of foraging rates

Here we present additional details for the field observations of foraging rates shown in Fig. S2.

Panel A of Fig. S2 shows the foraging rates for Colony 863 on September 5, 2015. The rates increase before reaching a QSS at around 10:30 am. The same colony on September 1, 2015 did not exhibit a QSS and had stopped foraging by 11 am (Fig. 4E). These observations are consistent with measurements showing that September 5, 2015 was a particularly cool and humid day while September 1, 2015 was much hotter and drier (see Table S1).

Panel B of Fig. S2 shows the data for Colony D19 on August 8, 2016 (from video recording) and provides an example of a very early cessation of foraging where both outgoing and incoming rates reached zero well before 11:00 am, similar to Colony 863 on September 1, 2015 (Fig. 4E). Both August 8, 2016 and September 1, 2015 were very hot and dry days (see Table S1).

Panel C of Fig. S2 show the data for Colony 859 on August 20, 2017 (from manual recording) and provides an example where the initial transient took a long time before ramping up. The initial transient for Colony 859 on August 20, 2017 remained at around 0.01 ants/sec from 10 am to 11:15 am before increasing to about 0.4 ants/sec by 12:30 pm. August 20, 2017 was a cool and humid day (see Table S1). Colonies might prefer different ranges of temperature and humidity; on cool and humid days, colonies that prefer warmer temperatures might forage at slightly later times of the day than colonies that prefer more cool temperatures.

Panel D of Fig. S2 show the data for Colony 1107 on August 16, 2017 (from manual recording) and provides a different example of a slow transient; it takes from 8 am to 10:30 am for the foraging rates to increase from 0.3 ants/sec to around 0.9 ants/sec. During this period the number of foragers outside the nest reaches almost 2000. In this case August 16, 2017 was not a particularly cool or humid day (see Table S1). The long transient and large number of active foragers suggests that the average time it took a forager to find a seed was long. Long foraging trip times can result in slow transients and high number of active foragers because when foragers take a long time to find a seed, it takes longer for foragers to return to the nest and interact with available foragers who then become active foragers. As well, when the average foraging trip times is long, more foragers might be required to cover larger and less dense foraging areas.

Panel E and F of Fig. S2 show the data for Colony 1017 on August 23, 2016 (from manual recording) and for Colony 1015 on August 18, 2016 and provide two examples of a burst in the outgoing foraging rate at the start of the foraging day that rapidly increases the number of active foragers outside the nest; it takes from 7:30 am to 7:45 am for Colony 1017 on August 23, 2016 to increase the number of active foragers from 0 to 800 and it takes from 7:15 am to 7:30 am for Colony 1015 on August 18, 2016 to increase the number of active foragers by 600. In both cases, the foraging rates reach a QSS that lasts tens of minutes. Both August 23 and August 18, 2016 were very dry days (see Table S1).

The burst kick starts the foraging process very rapidly and appears to be different from the mechanism that activates available foragers to leave the nest through interactions between incoming successful foragers and the available foragers. The rapid increase in the number of active foragers outside the nest might be advantageous on hot and dry days on which there will be only a short period of time in the early morning with acceptable foraging conditions.

## SI Appendix 3

### Analytical Approximation for 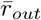 in terms of 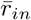 and c

Under the assumption that λ_*in*_ is a Poisson process with constant rate 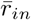, Eq. (4) is equivalent to a Poisson shot noise process with exponential decay:

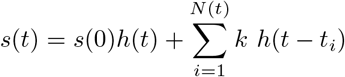

where *t*_*i*_ are the jump times of the Poisson process, and

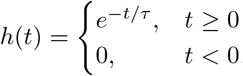

The mean and variance of this random process for an initial condition *s*(0) = 0 are given by 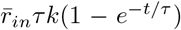 and 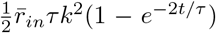 respectively [3]. Shot noise processes are Markovian and it can be shown that for finite jump sizes, *k* < ∞, *s* is ergodic [4], meaning that as *t* → ∞, *s*(*t*) converges in total variation to a unique stationary probability distribution π(*s*) for any initial condition *s*(0). In other words, *s* has the property that time averages converge in time to statistical averages. The ergodicity of *s* allows us to find an asymptotic expression as *t* → ∞ for the expected fraction of time that any single outcome of the random process spends in a region (*b*_1_,*b*_2_) by looking at its stationary probability density function.

Let *S*_*f*_ = {*t*_*f*_ ϵ [*t*_0_, *t*_0_ + *T*] | *b*_1_ < *s* < *b*_2_} be the set of all times over the time interval [*t*_0_,*t*_0_ + *T*] for which the stimulus is in the (*b*_1_,*b*_2_) region. Then *S*_*f*_ ⊆ *S* where *S* ={*t* ∈ [*t*_0_, *t*_0_ + *T*]}. We define 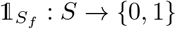 to be the indicator function associated with the subset *S*_*f*_

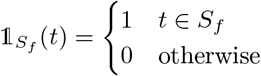

Let *T*_*f*_ be the amount of time that *s* is between *b*_1_ and *b*_2_:

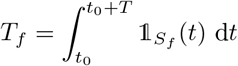

From the ergodic properties of *s*, and by the strong law of large numbers,

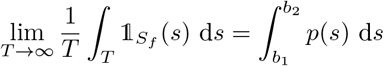

where *p*(*s*) is the density associated with *π*(*s*), i.e. the stationary probability density function (PDF) of *s*:

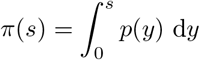

The PDF (see SI Appendix 4) is given as a piecewise function *p*_*i*_(*s*) for (*i* − 1)*k* ≤ *s* ≤ *ik* where the piecewise elements satisfy recurrence equations that depend on 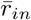, τ and *k*.

Let *b*_1_ and *b*_2_ be the FN bifurcation values of the input to the FN that takes the system from quiescence into the oscillatory regime and from this regime into saturation respectively:

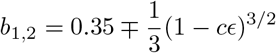

The size of the oscillatory region is given by the difference between *b*_2_ and *b*_1_ and it decreases with increasing volatility *c* (see SI Fig S4). For constant *s* where *b*_1_ ≤ *s* ≤ *b*_2_, the output rate is a constant given by the oscillation frequency of the FN when driven by a constant input *s*.

For *s* not constant, the FN transitions between quiescence, oscillatory behavior, and saturation as *s* varies. For *ϵ*_1_ ≪ 1, the FN dynamics are much faster than the dynamics of *s*, and the number of foragers leaving the nest in a given time period [*t*_0_,*t*_0_ + *T*] is proportional to *T*_*f*_, the amount of time spent by *s* in the oscillatory region.

For *T* → ∞, nonlinear effects in the oscillations become negligible and the mean outgoing rate becomes

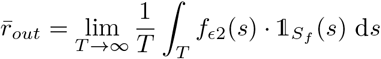

where *f*_*ϵ*2_ is the mean oscillation frequency of the FN when the driving input is constant and equal to *s*. We approximate *f*_*ϵ*2_ = 1/*T*_*LC*_ through the asymptotic representation [2]:

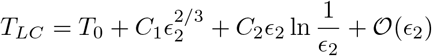

where *T*_0_, *C*_1_, and *C*_2_ are given in SI Appendix 1 to obtain an approximate expression for how 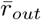 changes as a function of both 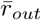 and *c*:

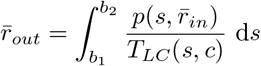

## SI Appendix 4

### Probability Density Function of s(t)

Here we find an analytical description of the probability density function of the stimulus function *s*(*t*) under the assumption that the incoming rate is a Poisson process. Under this assumption *s*(*t*) takes the form of a Poisson shot-noise process. Before we state our results, we state a result by Gilbert and Pollak. 1959 [5]:

#### Lemma 1.

*The amplitude distribution function F*_*s*_(*ξ*) = *Pr*[*s*(*t*)≤*ξ*] *for the shot noise process*

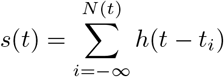

*where h(t) is called the impulse shape function, satisfies the integral equation*

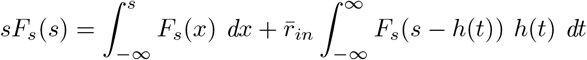

*Proof.* We refer the reader to [5] for the proof.

#### Theorem 2.

*The steady-state probability density function of the shot-noise process with exponential decay with impulses arriving with rate* 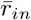

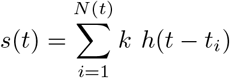

*where*

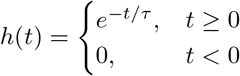

*is given as a piecewise function p*_*i*_(*s*) *for* (*i* − 1)*k* ≤ *s* < *ik where the piecewise elements satisfy the recurrence equations*:

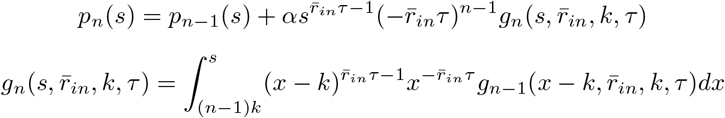

*with*

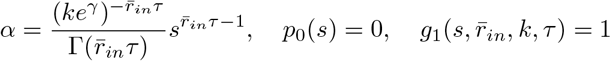

*where* γ = 0.5772… *is Euler’s constant and* Γ *is the gamma function.*

*Proof.* For a Poisson shot noise process in with impulse shape function:

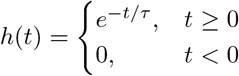

the integral equation in Lemma 1 can be rewritten as

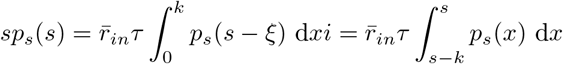

where *p*(*s*) = d*F*_s_/d*s* be the density function of *s*.

Differentiating with respect to *s*, we obtain

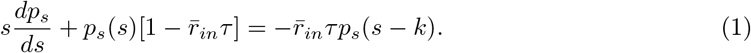

When 0 ≤ *s* ≤ *k*, *P*_*s*_(*s* − *k*) = 0, and

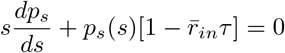

Picking 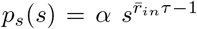 satisfies the equation above, so we have obtained a solution for *p*_*s*_(*s*) when 0 ≤ *s* ≤ *k*. For larger values of *s*, the differential equation Eq. (1) may be converted to an integral form

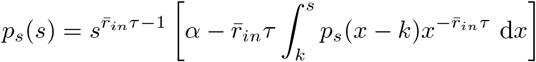

Since the integrand is known for *x* < 2*k*, we can determine *p*_*s*_(*s*) for *s* < 2*k*. This in turn enables us to integrate further to get *p*_*s*_(*s*) for *s* < 3*k*, etc. Let 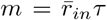, then the results for the first three jump regions *p*_*i*_, *i* = 1, 2, 3 are given by

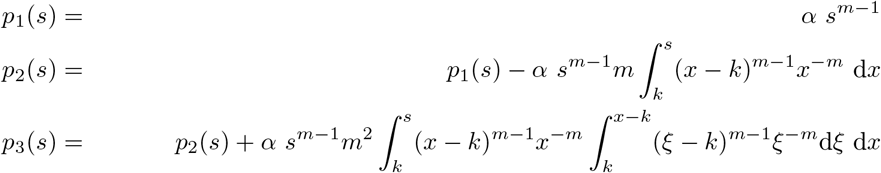

We now show by induction that *P*(*s*) satisfies the recurrence equations

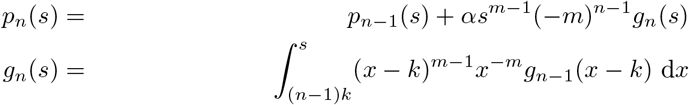

with *p*_0_ = 0, *g*_1_(*s*) = 1. For *n* = 1,

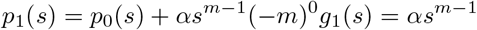

as expected. Now, assume that for *n* = *j*

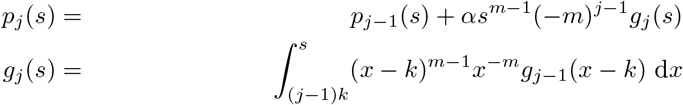

Then, for *n* = *j* + 1

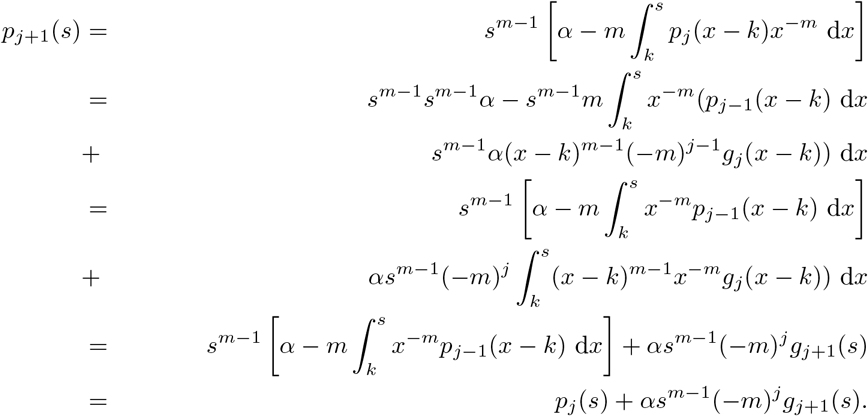

Finally, the constant α must be determined by the condition

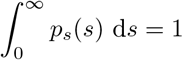

To compute the constant, we first note that the characteristic equation of *s* is given by

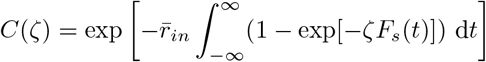

(see [3] for derivation). The characteristic function is the Laplace transform 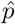 of *p*_*s*_,

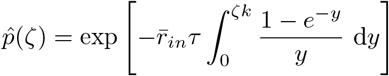

Using partial integration, this can be rewritten as

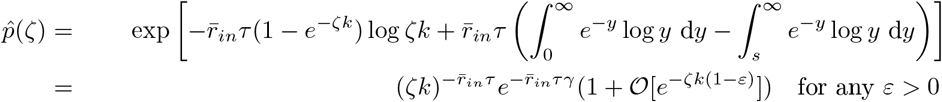

Thus, for 0 ≤ *s* ≤ *k*,

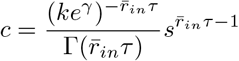

where γ = 0.5772… is Euler’s constant and Γ is the gamma function.

